# Microenvironmental TGF-β is an early driver of NF1-associated tumour formation

**DOI:** 10.1101/2025.06.23.661037

**Authors:** Alex Power, Salome Stierli, Elizabeth Harford-Wright, Guillem Modol-Caballero, Emma Lloyd, Stella Kouloulia, Giulia Casal, Lara Krasinska, Omar Bouricha, Ilaria Napoli, Sara Ribeiro, Cristina Venturini, Melanie P. Clements, Simona Parrinello, Alison C. Lloyd

## Abstract

Neurofibromatosis Type 1 (NF1) is a common tumour predisposition syndrome characterised by neurofibromas – *Nf1*^-/-^ Schwann cell (SC)-derived tumours of peripheral nerves. We and others have shown that *Nf1* loss in SCs is insufficient for neurofibroma formation but cooperates with an injury microenvironment to form tumours, but the mechanisms remained unknown. Here, we identify macrophage-secreted TGF-β as the microenvironmental injury signal that is essential for tumourigenesis. Analysis of the earliest stages of neurofibroma formation showed that tumours arise from a population of *Nf1*^-/-^ SCs that ‘escape’ the regenerating nerve shortly after injury. Here, they reside in a distinct microenvironment conducive for tumourigenesis, where TGF-β disrupts SC/axonal interactions and SC re-differentiation. Pharmacological inhibition of TGF-β for a short therapeutic window during this early stage inhibited tumour formation, highlighting the potential to normalise *Nf1*^-/-^ SCs and identifying TGF-β as a potential therapeutic target to both treat and prevent neurofibroma formation.

## Introduction

The common tumour predisposition disorder, Neurofibromatosis Type 1 (NF1), is characterised by the development of multiple benign cutaneous and plexiform neurofibromas^1^. These peripheral nerve-associated tumours cause considerable morbidity, pain, and disfigurement with a proportion of plexiform neurofibromas developing into malignant peripheral nerve sheath tumours (MPNST) that are refractory to treatment^2-5^. *NF1* encodes for the Ras-GAP neurofibromin, with loss of function resulting in hyperactive Ras signalling which has been shown to be responsible for tumourigenesis^6-9^. Schwann cells (SCs) or their precursors have been identified as the cell of origin of neurofibromas ^10,11^. Many of these models have also highlighted an important role for the microenvironment, with an injured environment either required or increasing the frequency of tumour development^12-17^.

Peripheral nerves are regenerative following an injury. This involves the de-differentiation of quiescent, adult SCs into proliferative ‘repair’ SCs that orchestrate the regrowth of damaged axons back to their targets^18-20^. This involves a complex multicellular response in which multiple cell types are coordinated by repair SCs to provide a conducive environment for axonal regrowth back to their targets. The de-differentiation of SCs to their repair state is driven by a sustained signal via the ERK-signalling pathway and it is considered that Nf1 loss primes SCs towards this de-differentiated state^10,21,22^. Neurofibromas have frequently been compared to unrepaired wounds and are composed of the same cell types that are involved in peripheral nerve regeneration, including ‘repair-like’ SCs and inflammatory cells associated with the repair process^2,10,17,23,24^. However, unlike the normal regenerative process which resolves, neurofibromas remain in this ‘repair’ state and continue to expand. Resolution of this injury response has been considered as a mechanism to prevent neurofibroma formation.

Our previous work established that *Nf1* loss in adult SCs does not result in neurofibroma formation, consistent with other studies^17,25^. However, following a nerve injury, Nf1^-/-^ SCs efficiently formed neurofibromas demonstrating that the microenvironment is key to determining whether Nf1-KO SCs will form tumours. Importantly, our work demonstrated that a specific niche microenvironment at the wound site was required for neurofibroma formation. Following a nerve transection, the two nerve stumps re-join to form new tissue at the wound site known as the bridge, that is initially composed of stromal and inflammatory cells. The regenerative process requires the regrowing axons to cross the bridge and then re-enter into the distal stump to grow back to their targets. Repair SCs migrate as cellular cords to guide regrowing axons across the bridge and also support the regrowing axons within the distal stump^18-20,26^. Once the axons reach their targets, the repair SCs re-differentiate to form new nerve tissue at the wound site and restore functionality within the distal stump. We showed that Nf1^-/-^ SCs only form neurofibromas at the site of injury (nerve bridge), with Nf1^-/-^ SCs distal to the injury behaving as normal SCs by re-associating with axons and re-differentiating^17^. This demonstrates that the distinct environments of the nerve bridge and the distal stump can determine whether an Nf1^-/-^ SC will undergo a tumour fate or revert to the behaviour of a normal SC. Importantly, by identifying that specific microenvironmental signals can determine whether tumours will form highlights the possibility of new therapeutic routes to both treat and prevent neurofibroma formation.

In this study, we sought to identify the microenvironmental factors responsible for Nf1^-/-^ SC tumour formation and characterise the mechanism by which normal nerve regeneration processes can give rise to a neurofibroma. Using our injury-induced neurofibroma model, we identified macrophage-derived transforming growth factor beta (TGF-β) as the microenvironmental factor that is required for tumour formation. We then show that the earliest stages of tumour formation involve the escape of Nf1^-/-^ SCs from the confines of the nerve environment, to form the mixed cell, disorganised structures, characteristic of neurofibromas, which is TGF-β dependent. Short-term TGF-β inhibition at this early stage was sufficient to impair neurofibroma formation providing promise for targeting this pathway therapeutically.

## Results

### TGF-β signalling is upregulated at the site of neurofibroma formation

We have previously shown that injury signals cooperate with Nf1 loss in SCs to induce neurofibroma formation in adult mice^17^. In this model, proliferating de-differentiated repair Nf1-KO SCs form neurofibromas at the injury site whereas Nf1-KO SCs revert to normal behaviour in the distal stump and re-differentiate in a manner indistinguishable to control cells (Figure 1A). To determine the nature of the signals responsible for these contrasting behaviours, we first identified the earliest timepoint at which the behaviour of Nf1-KO SCs diverged between the two nerve regions. To do this, we analysed the proliferation rates at early time points following nerve injury, comparing the bridge and distal stumps in Control (P0-CreER^T2^:YFP^fl/fl^:Nf1^+/+^) and Nf1-KO (P0-CreER^T2^:YFP^fl/fl^:Nf1^fl/fl^) mice, which additionally allowed the identification of recombined mSCs (YFP+). This analysis showed that at Day 7 following injury, Nf1-KO SCs proliferated at a higher rate compared to SCs in control animals in both the bridge and distal stumps demonstrating that Nf1-KO SCs initially have a proliferative advantage in both nerve regions. However, the proliferation rate of both Control and Nf1-KO YFP+ SCs was higher in the bridge region indicating that the bridge was a more proliferative environment than the distal stump (Figures 1B), consistent with our previous findings ^27^. By Day 10, the proliferation of Nf1-KO SCs in the distal stump had returned to the levels of Control SCs, whereas Nf1-KO SCs continued to proliferate at a higher rate in the bridge region, which was maintained at Day 14 (Figure 1B). Notably, the increased proliferation was limited to recombined SCs (YFP+), as unrecombined SCs (YFP-) in the nerve bridge proliferated at control levels, confirming that only Nf1-KO SCs co-operate with microenvironmental signals (Figure S1A-B). These results also suggested that very early events in the nerve repair process determine whether Nf1-KO SCs will form a tumour.

**Figure 1.**
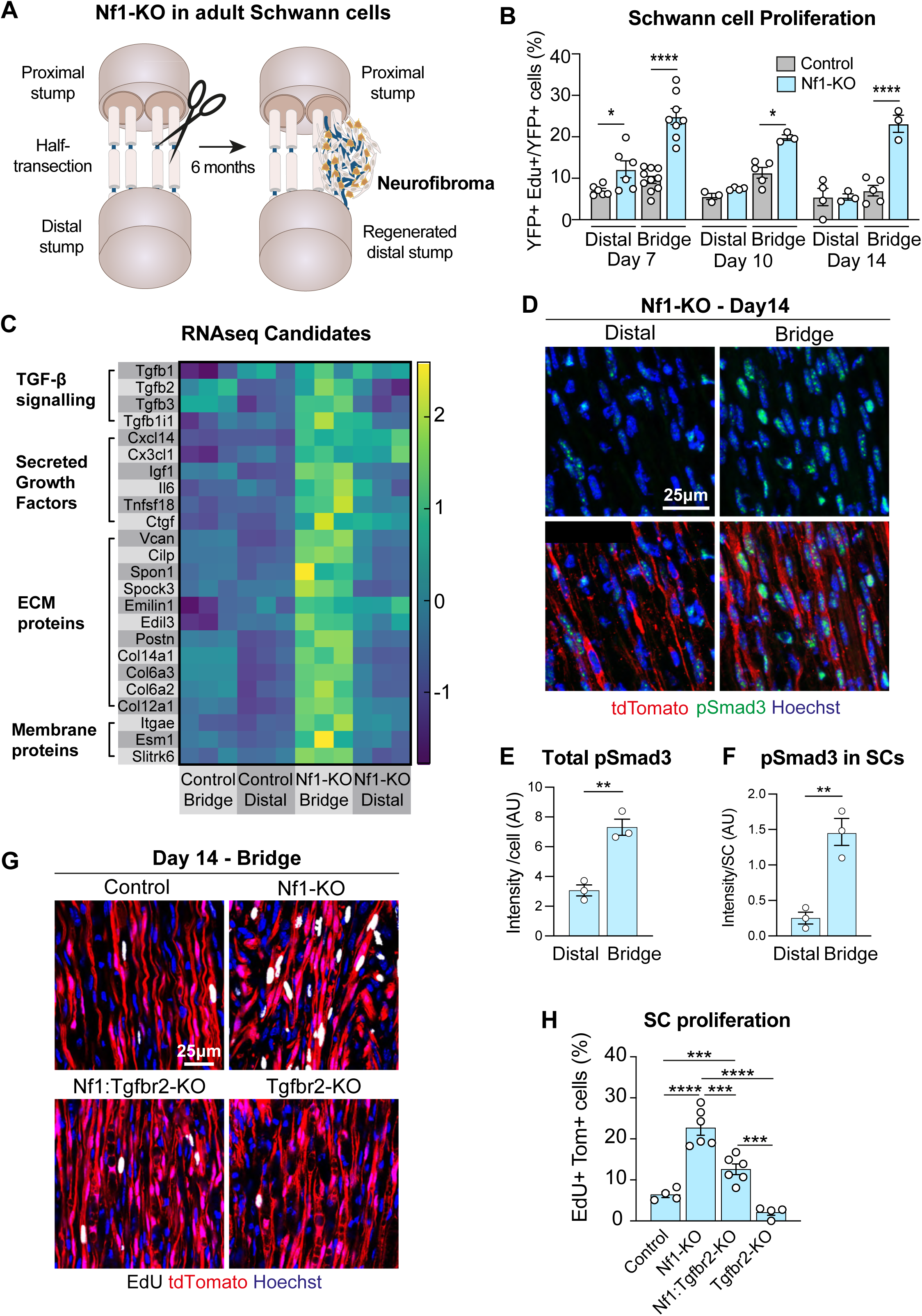
TGF-β signalling is upregulated at the site of neurofibroma formation. **A**. Schematic illustrating injury-induced Nf1-KO mouse model. Neurofibromas only form at the injury site within 6 months of injury. **B**. Quantification of percentage of EdU+/YFP+ SCs in the bridge and distal stumps at indicated days post-injury. Nerves were harvested 6h after EdU administration. Each dot represents one mouse (n=3-10), data shown as mean ± SEM. **C**. Heatmap depicting the z-scores of differentially expressed genes of interest from bulk RNA-seq analysis of the distal and bridge regions from Control and Nf1-KO animals at Day 14 post-injury. **D**. Representative confocal images of longitudinal sections of the bridge and distal stump of Nf1-KO mice at Day 14 post-injury, labelled to detect recombined SCs (tdTomato, red), p-Smad3 (green), and nuclei (Hoechst, blue). Quantification of (**D**) showing (**E**) pSmad3 intensity per cell or (**F**) per recombined SC. Each dot represents one mouse (n=3), data shown as mean ± SEM. **G**. Representative confocal images of longitudinal sections of Day 14 nerve bridges, showing recombined SCs (tdTomato, red), and labelled to detect EdU (white), and nuclei (Hoechst, blue). **H.** Quantification of (**G**), each dot represents one mouse (n=4-6), data shown as mean ± SEM. For B and H, a two-way ANOVA was used; for E and F, an unpaired two-tailed t-test was used. *p<0.05, **p<0.01, ***p<0.001, ****p<0.0001.

To take a non-biased approach to identify the microenvironmental signals responsible for tumour formation, we performed bulk RNA-seq analysis of the bridge and distal regions of Nf1-KO and Control mice at Day 14 after injury, the timepoint at which the proliferation differential was greatest between these regions. Principal component analysis showed separation of the groups (Figure S1C) and analysis of the transcripts showed many differentially regulated transcripts were specific to the wound site with a further group specific to Nf1-KO animals (Figures S1D). We speculated that secreted growth factors, ECM proteins, and membrane proteins were most likely to be involved and identified a number that were upregulated within the bridge region (Figure 1C). Amongst these candidates were TGF-β ligands, together with downstream targets of this pathway, such as versican and connective tissue growth factor^27-34^. We confirmed the increased transcript expression of a number of these candidates by RT-qPCR, validating the screen (Figure S1E). Increased TGF-β signalling in this region was further validated by the detection of higher levels of phosphorylated Smad3 (a canonical target of TGF-β) in SCs in the bridge region of Nf1-KO animals (Figures 1D-F)^35^. In previous work, we identified a role for TGF-β to maintain SCs in a more mesenchymal-like state, required for their migratory role transporting regrowing axons across the injury site^27^. Moreover, published expression data of human and mouse neurofibromas detected an increase in *Tgfb2* mRNA levels compared to control peripheral nerve ^36^, which we found was also increased in the Nf1-KO bridge at Day 14 post-injury (Figure S1F). Together these findings made the TGF-β pathway a strong candidate for further study.

To address whether TGF-β signalling in Nf1-KO SCs is important for neurofibroma formation, we generated mice in which TGF-β signalling was specifically ablated in recombined mSCs in Control and Nf1-KO mice. To do this, we crossed our P0-CreER^T2^:YFP^fl/fl^:Nf1^fl/fl^ mice^17^ to tdTomato^fl/fl^:Tgfbr2^fl/fl^ mice^27^, permitting the conditional knockout of the TGF-β ligand-binding receptor (TGFβR2), an obligatory component of the TGF-β receptor complex, specifically in mSCs^37^. The YFP reporter was also replaced with a tdTomato reporter (Figure S1F), as we found that tdTomato fluorescence was better maintained after fixation.

Initially we addressed whether TGF-β was responsible for the increased proliferation of Nf1-KO SCs in the bridge at the Day 14 timepoint. Analysis showed that loss of TGF-β signalling in Nf1-KO SCs resulted in a significant reduction in proliferation, although not completely to control levels (Figures 1G-H). These findings demonstrated that TGF-β signalling in Nf1-KO SCs was at least partially responsible for their increased proliferation at the earliest stages of tumour formation and identified TGF-β as a strong candidate for the microenvironmental signal specific to the bridge region that is required for neurofibroma formation.

### TGF-β signalling in Nf1-KO Schwann cells is required for neurofibroma formation

Based on these findings, we then addressed whether TGF-β signalling in SCs was required for neurofibroma formation. Our previous work showed that large neurofibromas were visible at the injury site in Nf1-KO mice 5 – 8 months following a sciatic nerve injury, while the nerves in control mice regenerated and appeared indistinguishable from uninjured nerve^17^. We therefore harvested nerves from the four groups of mice, 6 months after injury. Visual analysis of nerves prior to dissection, confirmed that all Control nerves appeared indistinguishable from uncut nerves (Figures 2A-B). In contrast, 100% (8/8) of nerves from injured Nf1-KO mice exhibited a large region of abnormal growth around the injury site, in some cases spreading along the nerve, which we previously showed was the result of a tumour with the pathology of a neurofibroma^17^. Remarkably, no visible tumours were observed in the Nf1:Tgfbr2-KO mice (n=9). This finding was confirmed by quantification of the size of the injury site in the four groups of animals (Figure 2C-D). These findings demonstrate that neurofibroma formation was entirely dependent on TGF-β signalling in Nf1-KO SCs.

**Figure 2.**
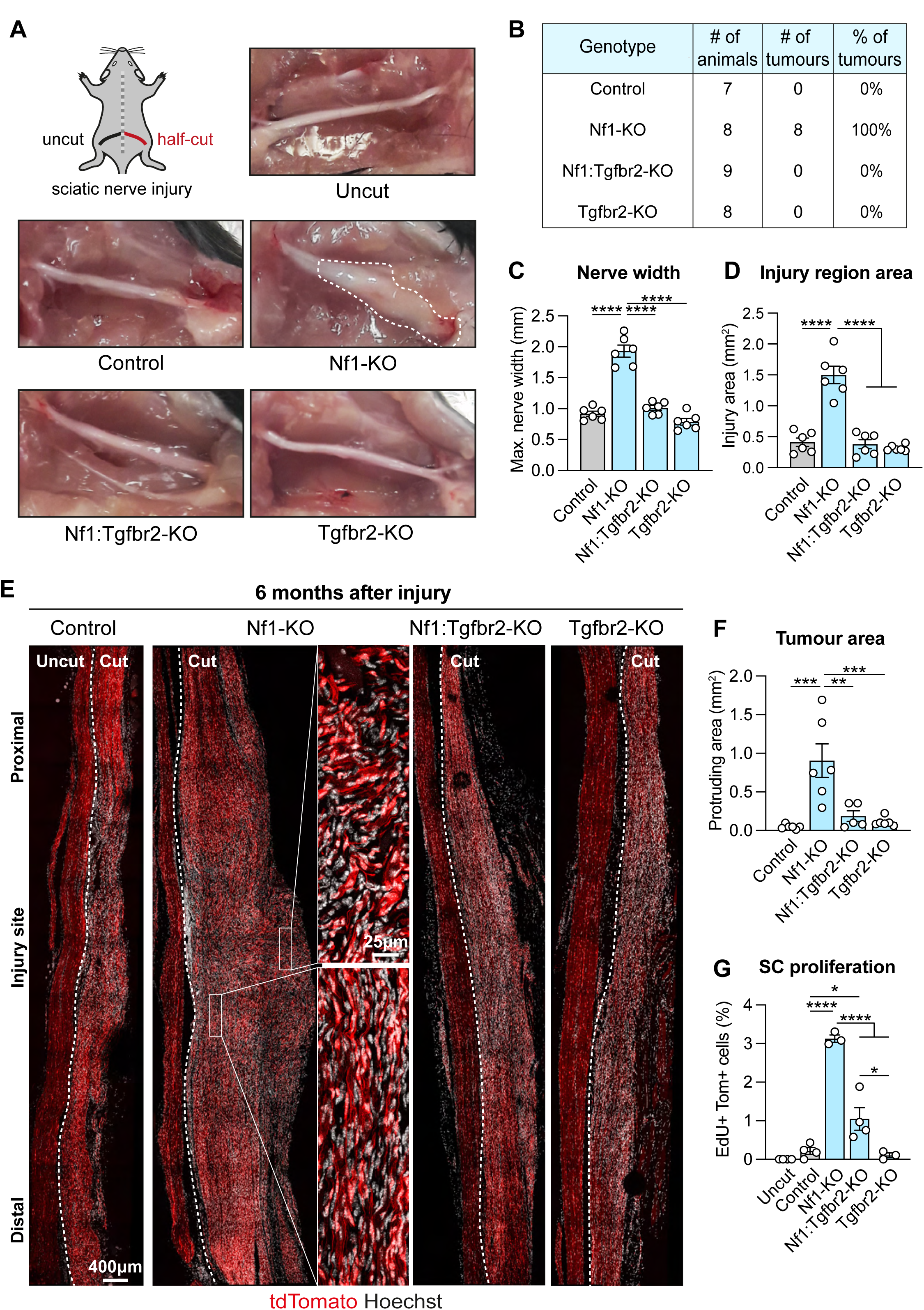
TGF-β signalling in Nf1-KO Schwann cells is required for neurofibroma formation. **A.** Schematic and representative macroscopic images of sciatic nerves 6 months after injury. The tumour is indicated by dashed lines. **B.** Table depicting the tumour frequency in indicated mice at 6 months post-injury. Quantification of (**A**) showing (**C**) maximum nerve width at the injury site and (**D**) injury region area. Each dot represents one mouse (n=6), data presented as mean ± SEM. **E.** Representative confocal tile scan images of longitudinal sections of injured nerves at 6 months after injury showing recombined SCs (tdTomato, red) and labelled to detect nuclei (Hoechst, white). Zooms show Nf1-KO tissue. Vertical dashed lines indicate the boundary between uncut and cut fascicles. **F.** Quantification of tumour area from (**E**). Each dot represents one mouse (n=5-6), data presented as mean ± SEM. **G.** Graph showing the percentage of proliferating tdTomato+ SCs at 6 months, determined by EdU labelling prior to harvest, related to images in **Figure S2C**. Each dot represents one mouse (n=3-4). Data presented as mean ± SEM. For C, D, F, and G, a two-way ANOVA was used. *p<0.05, **p<0.01, ***p<0.001, ****p<0.0001.

To analyse the structure of the tumours, we cut longitudinal sections of the nerves to visualise the proximal (upstream of the injury), bridge (injury site) and distal (downstream of the injury) regions of the regenerated fascicles, adjacent to uncut fascicles (Figure 2E). In all animals, the regenerated fascicle could be observed by virtue of increased Hoechst staining within the bridge region and along the distal stump, consistent with the increased cell density associated with a regenerated nerve^38^ (Figure S2A). In Control mice, despite the increase in cellularity, the regenerated region was a similar size to the region proximal to the injury, indicative of a normal regenerative response (Figures 2E). In contrast, and consistent with the visible tumours seen upon dissection, nerve sections from Nf1-KO mice exhibited a large bulge of cells emanating from the regenerated bridge region, many of which were tdTomato+, showing they derived from Nf1-KO SCs (Figures 2E-F and S2B). Reflecting the lack of visible tumours observed in Nf1:Tgfbr2-KO mice, a similar region was barely visible in nerve sections from these mice (Figure 2E-F). Proliferation analysis was consistent with these findings, in that tdTomato+ SC proliferation was higher in Nf1-KO mice and this was mostly dependent on TGF-β signalling (Figures 2G and S2C). Together these results show that the microenvironmental signal that is required for neurofibroma formation is TGF-β.

### Nf1-KO tumour tissue is distinct from Nf1-KO regenerated nerve

Closer observation of nerves from Nf1-KO mice showed that the bridge region was composed of two clearly distinct regions. An organised regenerated region reconnected to the distal stump and an adjacent tumour that appeared much more disorganised (Figures 3A-B). Consistent with this, Nf1-KO SCs in the regenerated region were associated with polarised axons and many expressed MBP, a marker of myelination (Figures 3C-E). Virtually all were also found within Glut1+ minifascicles (Figures 3C and 3F), that are characteristic of structures that form at the wound site to enclose regenerating SC/axonal structures^39^. Together these findings show that even within at the injury site, a proportion of Nf1-KO SCs do not form a tumour but instead re-differentiate to form a regenerated fascicle, displaying behaviour similar to SCs in Control animals.

**Figure 3.**
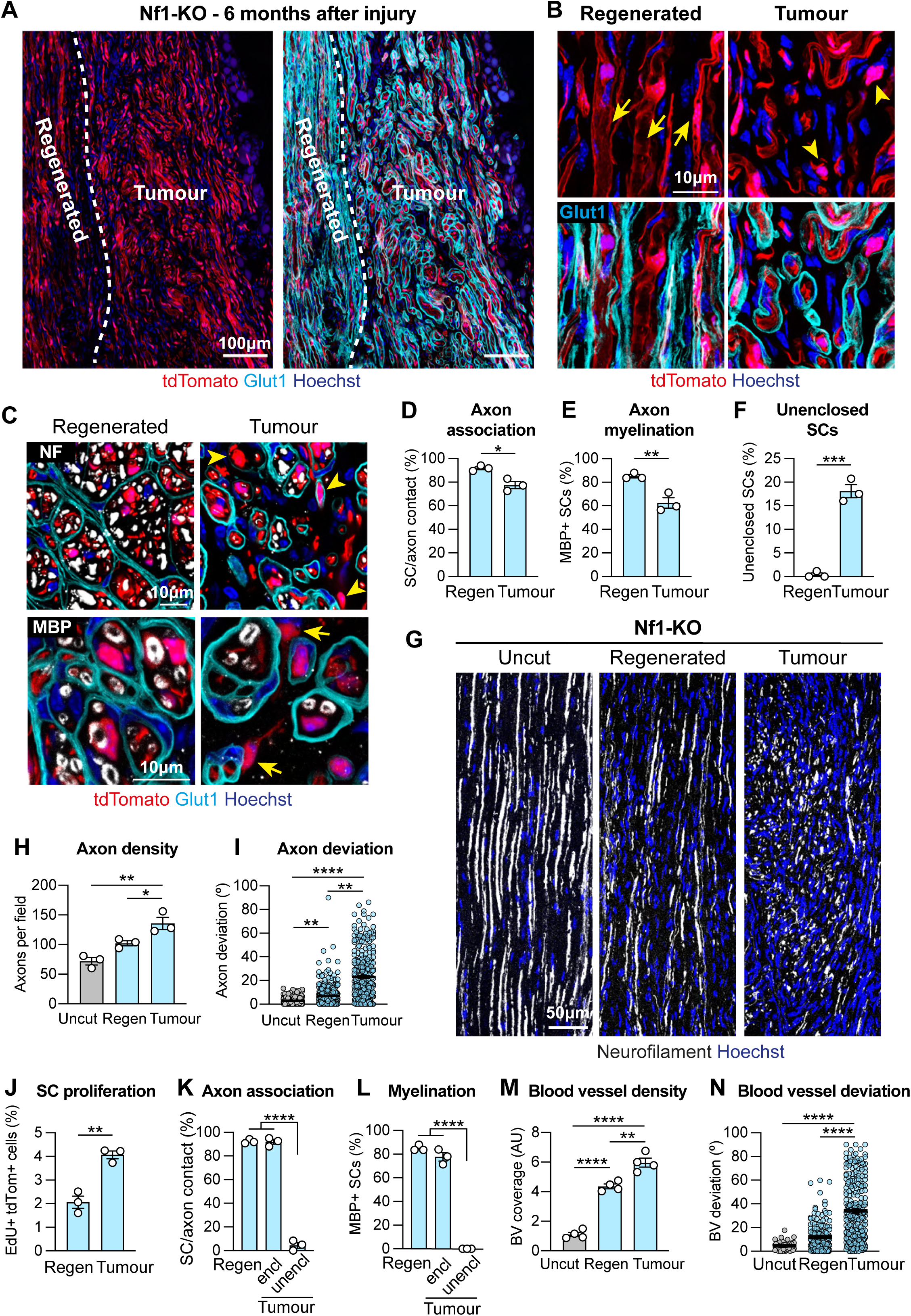
Nf1-KO tumour tissue is distinct from Nf1-KO regenerated nerve. **A.** Representative confocal images of longitudinal sections of Nf1-KO nerves at 6 months post-injury, showing recombined SCs (tdTomato, red), and labelled to detect perineurial cells (Glut1, cyan) and nuclei (Hoechst, blue). Dashed lines indicate the border between regenerated and tumour regions. **B.** Higher magnification images of (**A**). Arrows indicate recombined SCs with a differentiated morphology; arrowheads indicate abnormal SCs. **C.** Representative confocal images of transverse sections of regenerated and tumour regions of nerves from Nf1-KO mice at 6 months, showing recombined SCs (tdTomato,red), and labelled to detect Glut1 (cyan), myelin basic protein (MBP, white), neurofilament (NF, white), and nuclei (Hoechst, blue). Quantifications of (**C**) showing **(D)** the percentage of recombined SCs with axonal contact, **(E)** percentage myelination and **(F)** unenclosed by a Glut1+ perineurial minifascicle. Each dot indicates one mouse (n=3), data shown as mean ± SEM. **G.** Representative confocal images of longitudinal sections of uncut, regenerated and tumour tissue from Nf1-KO mice at 6 months post-injury, labelled to detect axons (NF, white) and nuclei (Hoechst, blue). Quantification of **(G)** showing **(H)** axonal density and **(I)** angle deviation from the proximal-distal axis. Each dot represents one mouse **(H)** or one axon **(I)**, data shown as mean ± SEM. **J.** Quantification of EdU+/tdTomato+SCs in the indicated regions, from NF1-KO mice, 6 months after injury. Each dot represents one mouse (n=3), data shown as mean ± SEM. Quantifications of (**C**) showing the percentage of recombined SCs, enclosed and unenclosed by minifascicles, that are **(K)** in axonal contact and **(L)** MBP+. Each dot represents one mouse, data shown as mean ± SEM. Quantification showing **(M)** blood vessel density and **(N)** angle deviation from the proximal-distal axis, in indicated regions of nerves from NF1-KO mice, 6 months after injury, related to **Figure S3B**. Each dot represents one mouse, data shown as mean ± SEM. For D-F, and J, an unpaired two-tailed t-test was used; for H, I, M, and N, a one-way ANOVA was used; for K and L, a two-way ANOVA was used. *p<0.05, **p<0.01, ***p<0.001, ****p<0.0001.

In contrast, Nf1-KO SCs within the tumour region appeared highly disorganised (Figures 3A-B). The tumours were innervated at a higher density than the regenerated region, but the axons were highly disorganised, mirroring the disorder of the SCs (Figures 3G-I and S3A). Moreover, consistent with a tumour phenotype, a lower proportion of re-differentiated mSCs (tdTomato+/MBP+) were found within the tumour region, and a higher number were found disassociated from axons (Figure 3C-E). Consistent with this failure to re-differentiate, we also observed more proliferation of Nf1-KO SCs in the tumour compared to the regenerated region (Figure 3J). Minifascicles were observed within the tumours, however these were disorganised compared to those in the adjacent regenerated region (Figures 3A-C). Interestingly, while nearly all Nf1-KO SCs in the regenerated region resided within a Glut1+ minifascicle, a population of Nf1-KO SCs in the tumour were found outside of these structures (Figures 3C and 3F). Remarkably, unenclosed Nf1-KO SCs interacted less frequently with axons, and all failed to re-differentiate (Figures 3C and 3K-L).

The tumour region displayed other known characteristics of neurofibromas. The tumours were densely vascularised with disorganised vessels (Figures 3M-N and S3B-C). Moreover, the tumours were infiltrated by inflammatory cells such as mast cells, macrophages, and T cells (Figures S3B-G), as previously reported^40,41^. Interestingly, while these properties were mostly enhanced in the tumour region compared to regenerated regions in Nf1-KO mice, the regenerated region from Nf1-KO mice exhibited higher levels of blood vessels and inflammatory cells than the regenerated regions of the other mouse groups (Figures S3A-G). This indicates that the regenerated fascicle in the Nf1-KO mice is not completely normal, retaining a more inflammatory profile that is mostly dependent on TGF-β signalling. Notably, Tertiary Lymphoid Structures, heavily enriched in both T cells and B cells, were found exclusively in Nf1-KO mice (Figure S3H). Together these results demonstrate a further demarcation of the microenvironment of the wound site into two distinct regions; with Nf1-KO SCs in one region forming a tumour, while others re-differentiate to contribute to the formation of a more organised, regenerated fascicle that resembles the normal regeneration process, albeit more highly vascularised and inflammatory (Figure S3I).

### Early tumourigenesis involves the formation of an escaped region from the regenerating nerve

To determine how the tumours form, we analysed the architecture of nerves at the earliest stages of tumour formation. As previously reported, perineurial cells (PnC) form minifascicles within the bridge region of a regenerating nerve, that can be used to demarcate the region of new nerve tissue formed within a regenerating nerve ^39,42^. Longitudinal sections of nerves from Control animals at Day 14, showed that the vast majority of tdTomato+ SCs in the nerve bridge were contained within the newly-formed PnC network (Figures 4A-C). In contrast, in Nf1-KO mice, a large area containing tdTomato+ SCs was observed outside of the confines of the regenerating fascicle (Figures 4A-C and S4A-B). We termed these ‘escaped’ cells due to their abnormal localisation. Notably, the formation of the ‘escaped’ region was dependent on TGF-β signalling, as it was mostly absent in Nf1:Tgfbr2-KO animals (Figures 4A-C). Based on location and the dependence on TGF-β signalling, we reasoned that the escaped region represented the earliest stages of tumour formation. Consistent with this, the region resembled the tumour in that it was highly disorganised compared to the regenerated region (Figures 4A, 4D and S4A-B). Importantly, Nf1-KO SCs in the escaped region were more proliferative than their regenerating region counterparts (Figures 4E-F), implying that the microenvironment of the escaped region fosters enhanced SC proliferation in a TGF-β-dependent manner.

**Figure 4.**
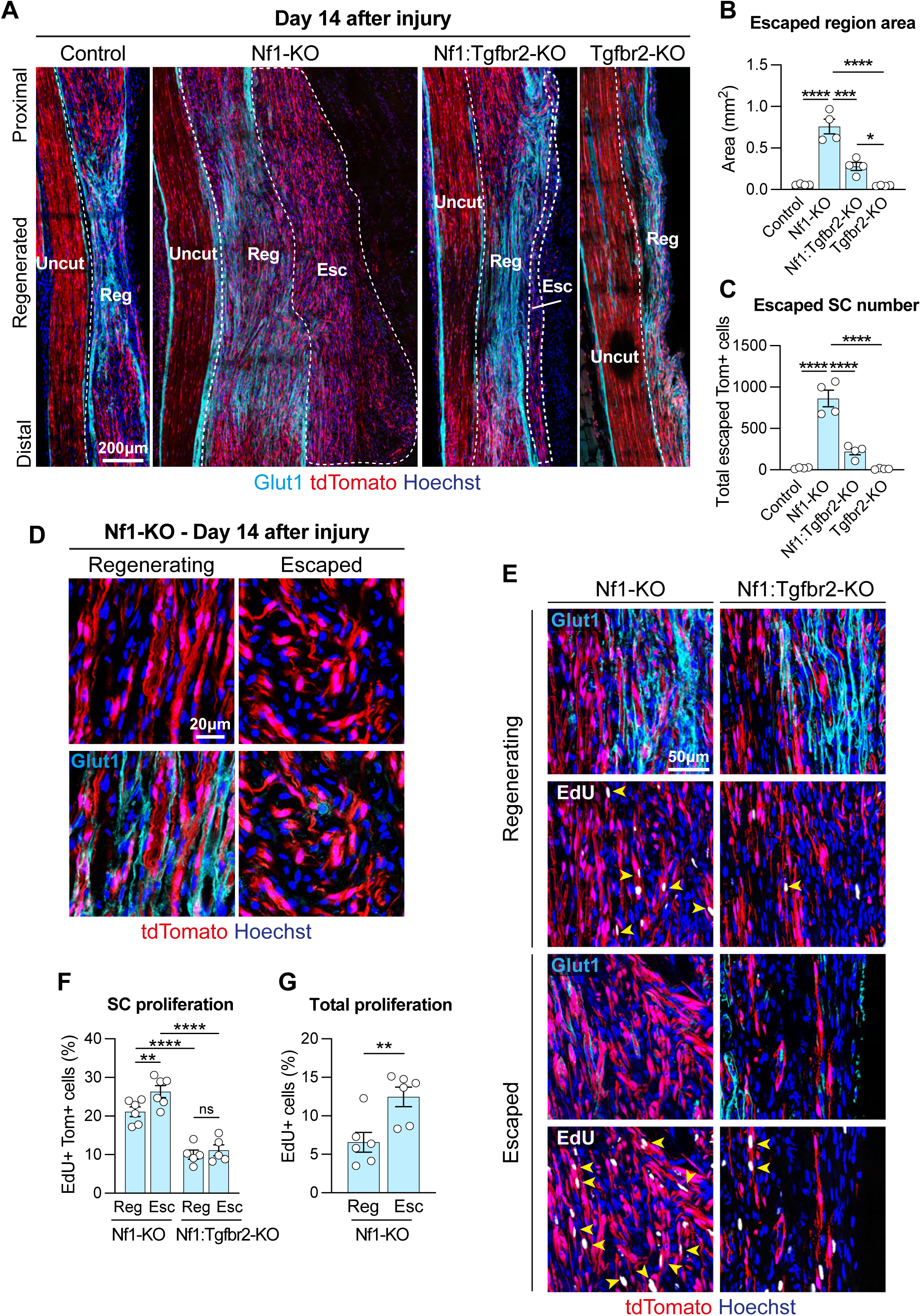
Early tumourigenesis involves the formation of an escaped region from the regenerating nerve. **A.** Representative tile scan images of longitudinal sections of nerves from indicated mice showing indicated regions at Day 14 after injury, showing recombined SCs (tdTomato, red) and labelled to detect perineurial cells (Glut1, cyan) and nuclei (Hoechst, blue). Dashed lines indicate the border between indicated regions. Quantifications of **(A)** showing (**B**) escaped region area and (**C**) number of escaped recombined SCs at Day 14 post-injury. Each dot represents one mouse (n=4), data shown as mean ± SEM. **D.** Representative confocal images of longitudinal sections of indicated regions of nerves from Nf1-KO mice at Day 14 post-injury, showing recombined SCs (tdTomato, red), and labelled to detect perineurial cells (Glut1, cyan) and nuclei (Hoechst, blue). **E**. Representative confocal images of longitudinal sections of indicated nerve regions at Day 14 post-injury in indicated mice, showing recombined SCs (tdTomato, red), and labelled to detect perineurial cells (Glut1, cyan), EdU (white), and nuclei (Hoechst, blue). EdU was injected 6h and 3h prior to harvest. Quantification of **(E)** showing proliferation of **(F)** recombined SCs and **(G)** all cells in the regenerating and escaped region of nerves from indicated mice, at Day 14 post-injury. Each dot represents one mouse (n=5-6), data shown as mean ± SEM. For B, C, and G, a two-way ANOVA was used; for F, an unpaired two-tailed t-test was used. *p<0.05, **p<0.01, ***p<0.001, ****p<0.0001, ns=not significant.

Nerve regeneration across the injury site is a complex, multicellular process, initiated by hypoxic macrophages, leading to the formation of blood vessels that provide a substrate for the migration of the SC cords that transport regrowing axons across the wound site^18,43^. Consistent with early tumour formation reproducing aspects of the wound repair process^10,20^, we observed large numbers of macrophages within both the regenerating and escaped region, with a higher density of macrophages found in the escaped region (Figures 5A-C and S5A). We also observed a high density of disorganised blood vessels within the escaped region consistent with our prior finding that blood vessels are required for SC migration during the regenerative process^18,43^ and reflecting the disorganisation seen in the mature tumour (Figures 5E-G).

**Figure 5.**
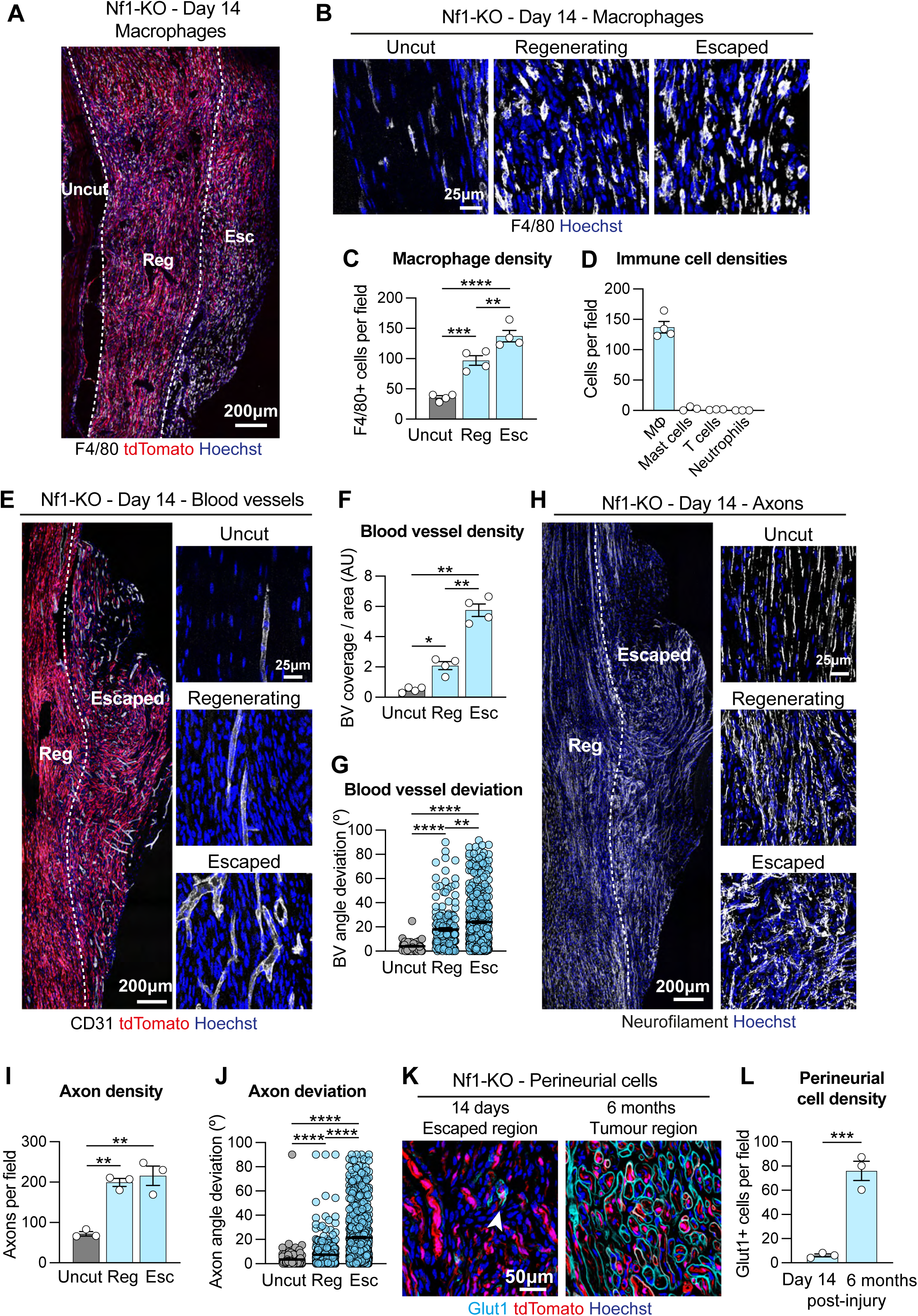
The escaped region is distinct from regenerated tissue. Representative tile scan confocal images (**A**) and zooms (**B**) of longitudinal sections of indicated regions from Nf1-KO mice at Day 14, showing recombined SCs (tdTomato, red), and labelled to detect macrophages (F4/80, white), and nuclei (Hoechst, blue). **C.** Quantification of (**A**) showing macrophage density. Each dot represents one mouse (n=4), data shown as mean ± SEM. **D**. Density of indicated immune cell-types at Day 14 in the escaped nerve region from Nf1-KO mice, related to **Figure S5B**. Each dot represents one mouse (n=3-4), data shown as mean ± SEM. **E.** Representative tile scan images and zooms of longitudinal sections of indicated nerve regions from Nf1-KO mice at Day 14 post injury, showing recombined SCs (tdTomato, red), and labelled to detect blood vessels (CD31, white), and nuclei (Hoechst, blue). Dashed lines indicate borders between the regions. Quantification of (**E**) showing (**F**) blood vessel density and (**G**) angle deviation from the proximal-distal axis. Each dot represents **(F)** one mouse (n=4) and **(G)** one blood vessel in. Data shown as mean ± SEM. **H.** Representative tile scan images and zooms of longitudinal sections of indicated nerve regions from Nf1-KO mice at Day 14 post injury labelled to detect axons (NF, white) and nuclei (Hoechst, blue). Dashed lines indicate the borders between the regions. Quantification of (**H**) showing (**I**) axon density or (**J**) angle deviation from the proximal-distal axis. Each dot represents (**I**) one mouse (n=3) or (**J**) one axon. Data shown as mean ± SEM. **K.** Representative confocal images of longitudinal sections of the escaped region at Day 14 and of the tumour region at 6 months post-injury from Nf1-KO mice, showing recombined SCs (tdTomato, red), and labelled to detect perineurial cells (Glut1, cyan), and nuclei (Hoechst, blue). Arrowhead indicates a rare Glut1+ cell in the escaped region. **L.** Quantification of (**K**) showing perineurial cell density. Each dot represents one mouse (n=3). Data shown as mean ± SEM. For C, F, G, I, and J, a one-way ANOVA was used. For L, an unpaired two-tailed t-test was used. *p<0.05, **p<0.01, ***p<0.001, ****p<0.0001.

As the nerve repair process proceeds, SCs respond to axonal signals and re-differentiate. We therefore reasoned that one possible explanation for the later expansion of the escaped region may be the absence of axons. However, we observed a similar increase in the number of axons in the escaped and regenerated regions, compared to the uncut nerve (Figures 5H-I). In contrast, similarly to the blood vessels and SCs, the axons were highly disorganised in the escaped region, consistent with a less directed, abnormal regenerative response (Figure 5J). Interestingly, in contrast to the tumours observed at 6 months, very few perineurial cells (Glut1+) (Figures 5K-L) or immune cells other than macrophages (mast cells, neutrophils or T cells) were found within the escaped region (Figures 5D and S5B), whereas they permeated the mature tumour (Figures S3B and S3E-I). This implies that both perineurial cells and these classes of immune cells do not have a role in the early stages of neurofibroma formation but may have a role in the establishment of the neurofibromas at a later stage.

In conclusion, the early stages of neurofibroma formation resemble the early stages of nerve regeneration, but this TGFβ dependent process is highly disorganised compared to the normal regenerative process and occurs outside of the reformed regenerating fascicle.

### Macrophages express TGF-β in the early tumour microenvironment

The formation of the escaped region is dependent on TGF-β signalling in Nf1-KO cells. To determine the levels and source of TGF-β in the escaped region, we used fluorescence *in situ* hybridisation (hybridisation chain reaction, HCR)^44^ to determine the expression of TGF-β mRNA in cell types within the different regions of the nerve. To confirm the specificity of this technique within an injured regenerating nerve, we first tested probes for *tdTomato* mRNA which should be restricted to recombined SCs. This showed a high level of specificity with little background staining (Figure 6A).

**Figure 6.**
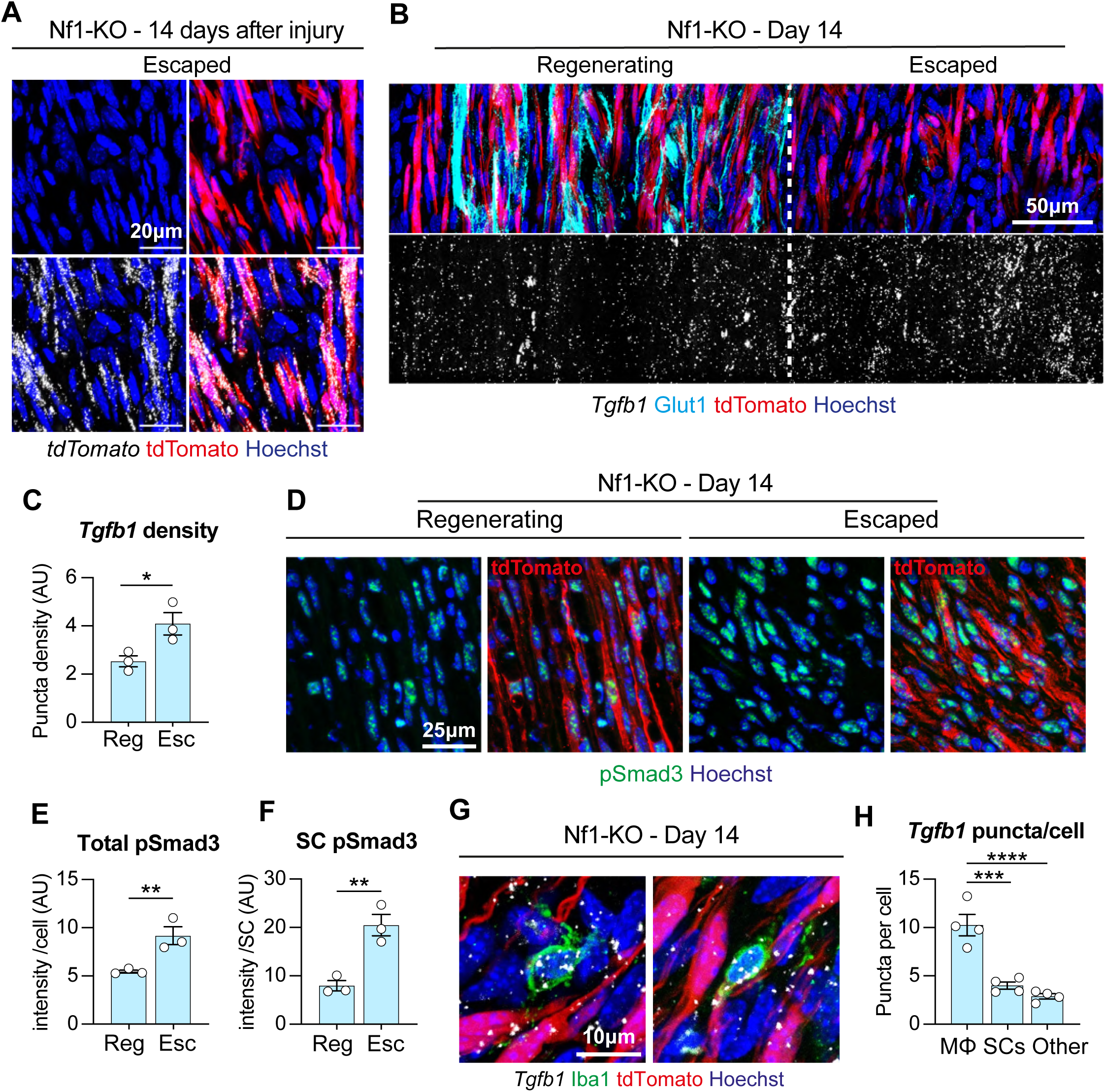
Macrophages express TGF-β in the early tumour microenvironment. **A.** Representative images of longitudinal sections of the escaped nerve region from NF1-KO mice at Day 14 post-injury, showing recombined SCs (tdTomato, red) and labelled to detect *tdTomato* mRNA (white) and nuclei (Hoechst, blue). **B.** Representative tile scan images of longitudinal sections of indicated nerve regions from Nf1-KO mice at Day 14 post-injury, showing recombined SCs (tdTomato, red) and labelled to detect *Tgfb1* mRNA (white), perineurial cells (Glut1, cyan), and nuclei (Hoechst, blue). **C.** Quantification of (**B**) showing the density of *Tgfb1* puncta in the indicated regions. Each dot represents one mouse (n=3), data shown as mean ± SEM. **D.** Representative confocal images of longitudinal sections of indicated regions from Nf1-KO mice, labelled to detect pSmad3 (green), recombined SCs (tdTomato, red) and nuclei (Hoechst, blue). Quantification of (**D**) showing (**E**) pSmad3 intensity per cell or (**F**) per recombined SC. Each dot represents one mouse (n=3), data shown as mean ± SEM. **G.** Representative confocal images of sections of the Nf1-KO escaped region at Day 14 post-injury, showing recombined SCs (tdTomato, red), and labelled to detect *Tgfb1* mRNA (white), macrophages (Iba1, green), and nuclei (Hoechst, blue). **H**. Quantification of (**G**) showing the number of *Tgfb1* puncta per indicated cell type. Each dot represents one mouse (n=4), data shown as mean ± SEM. For C, E, and F, an unpaired two-tailed t-test was used; for H, a one-way ANOVA was used. *p<0.05, **p<0.01, ***p<0.001, ****p<0.0001.

Analysis of *Tgfb* mRNA showed a gradient of expression from the regenerated to the escaped region with the highest level observed in the escaped region (Figures 6B**-**C). Consistent with this gradient of expression, we found higher levels of canonical TGF-β signalling activation, as measured by p-Smad3 levels, in cells, including SCs, within the escaped region (Figures 6D-F and Figure S6A). Together these results indicate that TGF-β signalling is increased in the escaped region at this early stage of tumour formation.

To identify the cell type responsible for TGF-β expression, we co-stained for markers of cells within the escaped region and found that macrophages (Iba1+) expressed a higher level of TGF-β mRNA compared to other cell types within the escaped region (Figures 6G-H). Together with our findings of an increased density of macrophages in the escaped region (Figures 5C and S6B), this indicates that macrophages are the major source of TGF-β in the developing tumour.

### TGF-β disrupts Schwann cell/axonal interactions and prevents Schwann cell differentiation

We have shown that in the early stages of tumour formation, Nf1-KO SCs escape from the fascicle and proliferate in a TGF-β-dependent manner. This continued proliferation could either be the result of TGF-β acting directly as a mitogen, or indirectly, by keeping SCs in a less differentiated, proliferative state. Support for the latter hypothesis came from our previous work, which showed that TGF-β maintained SCs in a more ‘mesenchymal’ state as they migrated across the bridge during normal nerve regeneration^27^. These studies also failed to find a role for TGF-β as a SC mitogen *in vivo* or *in vitro,* and consistent with these findings, we found that TGF-β exerted only a weak mitogenic effect on SCs alone or in combination with other factors (Figure S7A).

SC differentiation to a myelinating SC is a complex process requiring close association with axons, the establishment of a basement membrane, SC polarisation, followed by initiation of the Krox20-dependent myelination programme^45^. In contrast, differentiation to a non-myelinating SC is much less well understood but also requires a close association with axons to withdraw from the cell cycle^46,47^. To determine whether TGF-β disrupts SC/axonal interactions, we tested whether TGF-β affected the ability of SCs to interact with adult sensory neurons. Previously, we have shown that Nf1-knockdown disrupts the ability of SCs to interact normally with axons in a ERK pathway-dependent manner^46^, consistent with findings from some mouse models which identify disruption to SC/axonal interactions as the earliest observable event in neurofibroma formation^10,48,49^. Remarkably, we observed that TGF-β also disrupted SC/axonal interactions, but to a greater extent than Nf1 loss, and cooperated with Nf1 loss causing further disruption to axonal interactions (Figures 7A-B and S7B). The disruption by TGF-β was independent of the ERK-signalling pathway (Figures 7C-D), which is responsible for Nf1 effects^46^, but instead was dependent on signalling through the canonical Smad signalling pathway (Figures 7C-D). Notably, while TGF-β was able to disrupt the formation of normal SC/axonal interactions it failed to disrupt established interactions (Figures S7C-D), suggesting that TGF-β only exerts its effect in a ‘injury’ situation rather than having effects on stable nerves.

**Figure 7.**
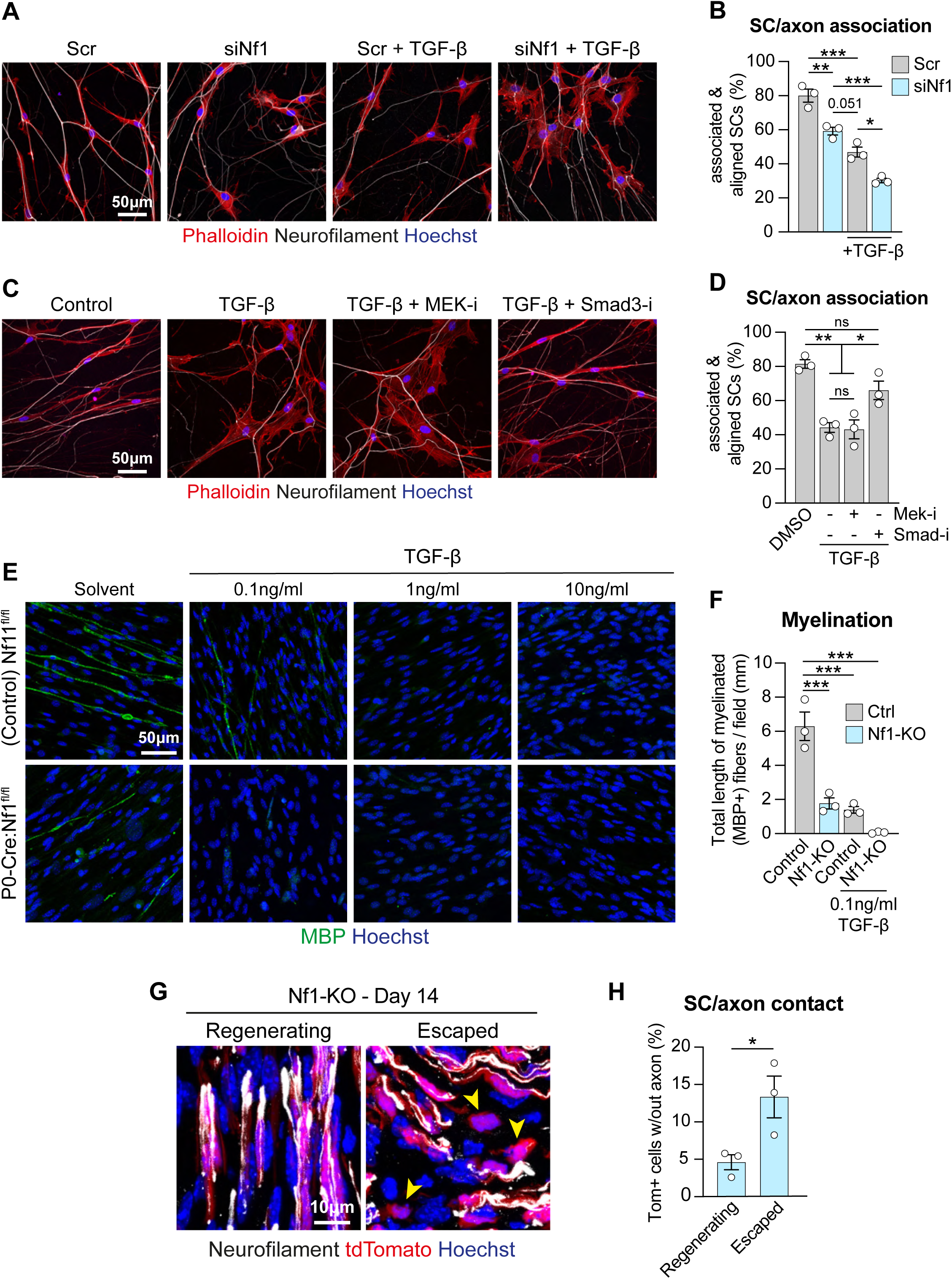
TGF-β disrupts Schwann cell/axonal interactions and prevents Schwann cell differentiation. **A.** Representative confocal images of DRG/SC co-cultures 12h after SC seeding, cultured in the presence or absence of 10ng/ml recombinant TGF-β. SCs were transfected with 2nM scrambled (scr) or Nf1-targeting siRNA, 48h prior to seeding. Cultures were labelled to detect axons (neurofilament, white), actin (phalloidin-546, red) and nuclei (Hoechst, blue). **B**. Quantification of (**A**) showing the percentage of SCs that had successfully associated and aligned to an axon. Data shown as mean ± SEM of n=3 independent experiments. **C.** Representative confocal images of DRG/SC co-cultures, 12h after SC seeding, cultured in the presence or absence of 10ng/ml TGF-β, MEK inhibitor PD184352 (0.5µM), or SMAD3 inhibitor SIS_3_ (5µM) as indicated. Cultures were labelled to detect axons (neurofilament, white), actin (phalloidin-546, red) and nuclei (Hoechst, blue). **D.** Quantification of (**C**) showing the percentage of SCs that had successfully associated and aligned to an axon. Data shown as mean ± SEM of n=3 independent experiments. **E.** Representative confocal images of DRG/SC myelinating cultures harvested from Nf1^fl/fl^ (control) or P0-Cre:Nf1^fl/fl^ adult mice. Cultures were labelled to detect myelin (MBP, green) and nuclei (Hoechst, blue). **F.** Quantification of (**E**) showing the total length of MBP+ fibers per field of view. Each dot represents the average of at least 6 fields from cultures from an individual mouse. Data shown as mean ± SEM of n=3 independent experiments. **G.** Representative confocal images of longitudinal sections of nerves from Nf1-KO mice, showing recombined SCs (tdTomato, red), and labelled to detect axons (NF, white), and nuclei (Hoechst, blue). **H.** Quantification of (**G**) showing the percentage of recombined SCs without detectable contact with an axon. Data shown as mean ± SEM. For B, D, and F, a two-way ANOVA was used; for H, an unpaired two-tailed t-test was used. *p<0.05, **p<0.01, ***p<0.001, ns=not significant.

To determine whether TGF-β also affects SC differentiation, we used (i) a simple cell-based differentiation assay and (ii) myelination assays in adult DRG explants isolated from mice in which Nf1 is deleted from all SCs (P0-Cre:Nf1^fl/fl^)^48^. In both assays, TGF-β blocked SC differentiation (Figures 7E-F and S7E-F), with the effect independent of ERK signalling (Figure S7E). In the DRG myelination assays, we found that Nf1-KO SCs retained the ability to myelinate, consistent with *in vivo* behaviour^17,48,50^. However, the myelination was significantly less efficient than control SCs (Figures 7E-F) consistent with previous *in vitro* findings^51,52^. Interesting, the residual myelination was inhibited by a lower concentration of TGF-β, indicating that Nf1-deficient SCs exhibit increased sensitivity to TGF-β signalling (Figures 7E-F).

Together these results demonstrate that TGF-β disrupts SC differentiation via two separate but linked events. Firstly, by disrupting normal SC/axonal interactions and secondly, by blocking differentiation. Consistent with these findings, we observed that Nf1-KO SCs in the escaped region of regenerating nerves at Day 14 post-injury frequently exhibited aberrant interactions with axons (Figures 7G-H) and many failed to later re-differentiate as the tumour progressed (Figures 3C-E).

### Pharmacological TGF-β inhibition for a brief therapeutic window impairs neurofibroma formation

Our results demonstrate that early tumour development relies on a TGF-β dependent expansion of Nf1-KO SCs that escape from the regenerating fascicle. In contrast, those that remain in the fascicle, re-differentiate and behave more normally, indicating that Nf1-KO SCs are on the cusp of normal behaviour in a manner strictly dependent on the microenvironment. We therefore reasoned that treatment with a TGF-β signalling inhibitor during a brief therapeutic window might be sufficient to ‘normalise’ Nf1-KO SC behaviour and block subsequent tumour formation. We chose to use Galunisertib, a potent, specific TGF-β receptor inhibitor, that has been frequently used in mouse studies^53,54^.

We initially confirmed that the inhibitor blocked the effects of TGF-β in our *in vitro* differentiation assay (Figures S8A-C). We then tested whether a short treatment with the inhibitor was sufficient to inhibit the TGF-β-dependent proliferation seen at the early stages of tumour formation. Based on previous mouse studies^53^ and our findings, we chose to treat mice with 75mg/ml Galunisertib twice daily for a short treatment window between Days 7-9 following injury. This time window was chosen, as by Day 7, SCs and regrowing axons have largely traversed the nerve bridge, minimising potential effects on the regeneration process itself^27^. Strikingly, we found that nerves from Galunisertib-treated mice harvested at Day 14 exhibited a significant decrease in the size of the escaped region, and in the number, density, and proliferation of escaped SCs, very similar to the extent of inhibition seen in the genetic KOs (Figures 8A-C and S8D-F).

**Figure 8.**
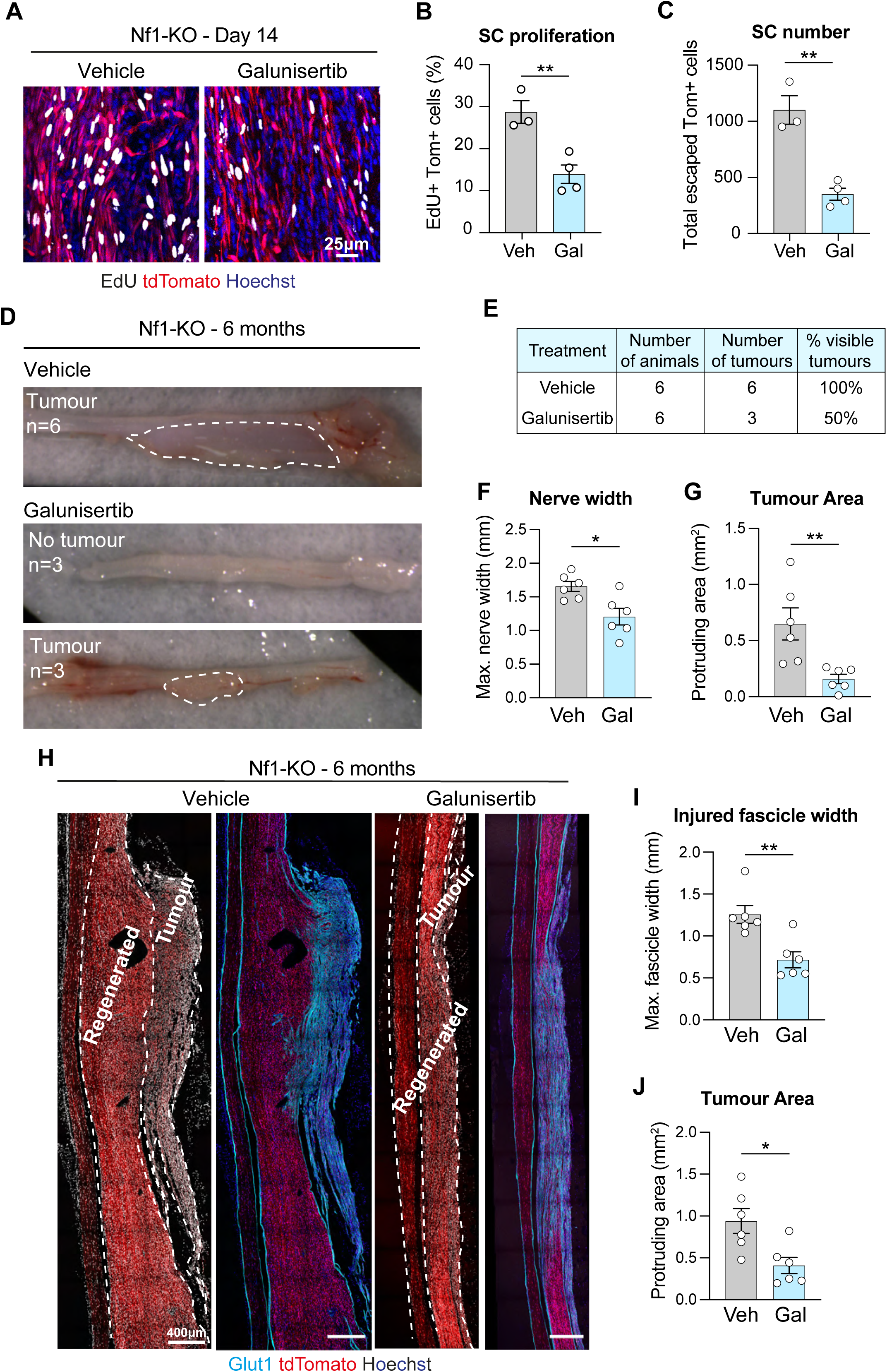
Pharmacological TGF-β inhibition for a brief therapeutic window impairs neurofibroma formation. **A.** Representative confocal images of longitudinal sections of nerves from Vehicle or Galunisertib-treated Nf1-KO mice at Day 14 post-injury, showing recombined SCs (tdTomato, red), and labelled to detect EdU (white) and nuclei (Hoechst, blue). EdU was injected 6h and 3h prior to harvest. **B.** Quantification of (**A**) showing SC proliferation. Each dot represents one mouse, data shown as mean ± SEM. **C.** Quantification of the number of escaped SCs at Day 14 post-injury in Vehicle and Galunisertib-treated Nf1-KO mice, relating to **S8D**. Each dot represents one mouse, data shown as mean ± SEM. **D.** Representative macroscopic images of Nf1-KO nerves at 6 months post-injury following treatment with Vehicle or Galunisertib. Dashed lines indicate tumour area. **E.** Table of tumour frequency. Quantification of (**D**) showing (**F**) maximum nerve width at the injury site and (**G**) tumour area in Vehicle and Galunisertib-treated animals at 6 months. Each dot represents one mouse (n=6), data shown as mean ± SEM. **H.** Representative tile scan images of longitudinal sections of nerves from NF1-KO mice, treated with Vehicle or Galunisertib and harvested at 6 months, showing recombined SCs (tdTomato, red), and labelled to detect perineurial cells (Glut1, cyan), and nuclei (Hoechst, white). Quantification of (**H**) showing (**I**) the maximum width of the injured fascicle and (**J**) tumour area in Vehicle and Galunisertib-treated animals harvested at 6 months. Each dot represents one mouse (n=6), data is presented as mean ± SEM. For B, C, F, G, I, and J, an unpaired two-tailed t-test was used. *p<0.05, **p<0.01.

To determine whether this was sufficient to block tumour formation, we repeated the schedule and harvested nerves, six months following nerve injury. As expected, we found that 100% of vehicle-treated Nf1-KO mice developed visible tumours. In contrast, only 50% of Galunisertib-treated mice developed visible tumours and those with visible tumours were mostly smaller than in control animals (Figure 8D-G). Consistent with these observations, longitudinal sections showed the near absence of tumour in the majority of the treated mice (Figure 8H-J). Together these findings demonstrate that pharmacological inhibition of TGF-β signalling for a short time-window during early tumourigenesis was sufficient to inhibit neurofibroma formation, a finding with potentially significant clinical implications.

## Discussion

Neurofibroma formation, as for most tumour types, is influenced by the microenvironment^24^. Most compellingly, an injured microenvironment has been associated with enhanced neurofibroma formation in several different mouse models^12-16^. In our injury-associated mouse model^17^, the unique environment of the wound site was shown to be critical. In this model, adult Nf1-KO SCs only formed tumours at the site of injury to a nerve. In contrast, Nf1-KO SCs in the distal stump failed to form tumours, despite these cells de-differentiating and proliferating following the injury in an environment rich in inflammatory cells^17^. This model thus provided a powerful model system to explore both the microenvironmental signals responsible for promoting neurofibroma formation and also allowed the tracking of the early development of these tumours. Taking this approach, we have made several important findings. Firstly, that TGF-β is the microenvironmental signal responsible for neurofibroma formation, and that it acts directly on Nf1-KO SCs. Secondly, that neurofibroma formation involves a disruption to the normal regeneration process, in that the tumours arise from Nf1-KO SCs that escape from the reforming nerve fascicle, and thirdly, that within this escaped environment, TGF-β acts to block SC normalisation by both disrupting normal axonal interactions and re-differentiation.

By studying the earliest stages of tumour formation, we were able to observe that the tumours appeared to arise from a population of Nf1-KO SCs that ‘escape’ from the reforming, regenerating fascicle. A key question therefore is how this escaped region forms? During normal nerve regeneration following a transection injury, a multicellular process organises and guides the migrating Schwann cell cords, which transport regrowing axons across the injury site^18,43,55,56^. The escaped region therefore indicates a failure of the directionality of this process, as Nf1-KO SCs appear to guide axons in a disorganised manner, away from their normal trajectory. SCs themselves are guided by a polarised vasculature which forms in response to VEGF signalling from hypoxic macrophages^43^. It is noticeable that the escaped region is rich in macrophages and blood vessels, which are also highly disorganised. This supports the hypothesis that the escaped region represents an aberrant recapitulation of the normal repair process but why the directionality of the process is lost remains unclear.

Once escaped, Nf1-KO SCs continue to expand resulting in frank neurofibroma formation that is dependent on TGF-β signalling. We showed previously that TGF-β signalling was required for normal nerve repair by keeping SCs in a more mesenchymal state^27^ and we show here that TGF-β disrupts the ability of SCs to interact normally with axons and differentiate and that Nf1-KO SCs are more sensitive to these effects. Perhaps during normal repair, TGF-β keeps SCs in a more dedifferentiated state, so they do not respond to axonal differentiation signals until the axons have re-entered into the distal stump. In contrast, it appears that TGF-β levels and signalling remain higher in the escaped region, resulting in the wounds that fails to repair, as neurofibromas have been previously characterised^10^.

What is particularly striking about our findings is how readily Nf1-KO SCs can be enticed to behave normally. Previous work in many model systems have already shown that Nf1-KO SCs mostly develop normally^17,48,50^. What is revelatory about our findings, is that even at the site of tumour formation, Nf1-KO SCs in an adjacent region regenerate mostly normally and even within the tumour, a proportion of Nf1-KO SCs associated with axons and re-differentiated. This highlights the critical role of the very local microenvironment and also how readily Nf1-KO SCs can be persuaded to normalise. This was perhaps most clearly demonstrated by our findings that a short therapeutic treatment to block TGF-β signalling at the earliest stages of tumour formation, was sufficient to block or inhibit tumour development. This further suggests that tilting the balance towards differentiation would be a successful approach to both prevent and treat neurofibroma formation.

Interestingly as the tumours evolve, their composition and structure changes. Perineurial cells invade the tumour and appear to organise the SC/axonal bundles into structures resembling the minifascicles seen in the regenerated bridge region. Notably, many Nf1-KO SCs differentiate within these structures, whereas those outside fail to differentiate. This suggests that over time, the growing tumour may progress towards normalisation, which might be predicted to limit the size of the tumours - consistent with clinical observations that neurofibromas have a growing and then stationary phase ^57^. The tumours also develop a more inflammatory phenotype, with mast cells, T cells and neutrophils associated with the mature tumour, possibly reflecting a suppressive immune response that may limit tumour expansion. Together our results identify TGF-β as a microenvironmental signal essential for neurofibroma formation, acting to support the expansion of Nf1-KO SCs that escape the normal environment of the nerve by disrupting normal axonal interactions and differentiation. Moreover, our findings indicate that Nf1-KO SCs can be “normalised” by even a transient inhibition of TGF-β signalling, providing a new therapeutic approach for potentially treating and preventing neurofibroma formation.

## Resource Availability

### Lead contact

Further information and requests for resources and reagents should be directed to and will be fulfilled by the lead contact, Alison Lloyd.

### Materials availability

This study did not generate new unique reagents.

### Data and code availability

All data reported in this paper will be shared by the lead contact upon request. Any additional information required to re-analyse the data reported in this study is available from the lead contact upon request.

The bulk RNA-seq data have been deposited at GEO: GSE286371 and will be publicly available from the date of publication.

## Acknowledgments

This work was supported by a programme grant from Cancer Research UK (C378/A4308) and supported by a Sub-agreement from the Johns Hopkins University via the Neurofibromatosis Therapeutic Acceleration Program (NTAP) with funds provided by Grant Agreement from Bloomberg Philanthropies. Its contents are solely the responsibility of the authors and do not necessarily represent the official views of the Johns Hopkins University. We would like to thank UCL Biological Services for animal husbandry and useful advice, and the Lloyd lab for advice, assistance with surgeries and useful discussions.

## Author contributions

A.C.L. conceived the project. A.P. and S.S. performed the majority of experiments with help from E.H-W., G.M.C., E.L., S.K., G.C., L.K., O.B., I.N., S.R., C.V.,and M.P.C. E.H-W, and S.P. advised on experimental design and A.C.L. supervised the study. A.P., S.S., E.H-W and A.C.L interpreted data and wrote the manuscript. All authors discussed the results and commented on the manuscript.

## Declaration of interests

The following patent has been submitted related to this application, PCT Application No. GB2024/051610.

## STAR METHODS

### Experimental Model Details

#### Animals

All animal work was performed in accordance with UK Home Office guidelines at the Central Biological Services Unit at University College London, UK. Mice were housed in a temperature (20-24°C) and humidity (45-65%) controlled vivarium on a 12h light/dark cycle with free access to food and water. Both male and female adult mice were used. To generate the P0-CreER^T2^:tdTomato^fl/fl^:Nf1^fl/fl^:Tgfbr2^fl/fl^ mice used in this study, P0-CreER^T2^:YFP^fl/fl^:Nf1^fl/fl^ mice ^17^ were crossed to the P0-Cre:tdTomato^fl/fl^:Tgfbr2^fl/fl^ mice ^27^. To generate the P0-Cre:Nf1^fl/fl^ mice used in this study, P0-Cre mice ^58^ were crossed with Nf1^fl/fl^ mice ^59^.

#### Primary cell cultures

Primary rat Schwann cells were isolated from the sciatic and brachial nerves of male and female Sprague-Dawley rats at postnatal Day 7 as previously described ^60^. In brief, sciatic and brachial nerves were extracted, desheathed, and digested using a mixture of collagenase and trypsin. Contaminating fibroblasts and macrophages were removed by negative immunopanning. Schwann cells were subcultured up to a maximum of 14 times and maintained at 37°C and 10% CO_2_.

### Method details

#### Administration of substances

To induce Cre-mediated recombination, 200mg/kg Tamoxifen (Sigma) was orally administered to 4–5 week-old mice daily for 5 consecutive days. Tamoxifen was dissolved in ethanol and diluted 10-fold in sunflower seed oil (Sigma) to a final concentration of 20mg/ml.

To assay proliferation, 2mg of 5-ethynyl-2-deoxyuridine (EdU, Sigma)/PBS (ThermoFisher) was injected by the intraperitoneal (IP) route, 6h and 3h prior to harvest at Day 14 post-injury, and 48h, 24h, and 4h prior to harvest at the 6-month post-injury timepoint. EdU incorporation was detected by immunofluorescence using the Click-iT EdU imaging kit (Invitrogen) according to the manufacturer’s instructions.

To pharmacologically inhibit TGF-β signalling, 75mg/kg Galunisertib (LY2157299, MedChemExpress) was orally administered to mice twice daily on Days 7–9 after injury. Galunisertib was dissolved in 5% DMSO in H_2_O.

#### Sciatic nerve injury

Mice underwent surgery 10 days after the final tamoxifen administration. Sciatic nerve partial transection was performed under aseptic conditions and under isoflurane anaesthetic. The right sciatic nerve was exposed and partially transected with scissors 1–2mm below the notch.

The wound was closed using surgical clips and the animal allowed to recover in a pre-heated chamber. Surgical clips were removed 14 days after injury.

#### Cell culture

Primary rat Schwann cells (SC) were cultured on 100μg/ml poly-L-lysine (PLL, Sigma)-coated dishes in SC medium consisting of low glucose (1g/l) Dulbecco’s Modified Eagle’s Medium (DMEM, Gibco) supplemented with 3% foetal bovine serum (FBS, BioSera), 1μM forskolin (Abcam), 4mM L-glutamine (Gibco), 100μg/ml kanamycin (Gibco), 800μg/ml gentamicin (Gibco,), and glial growth factor (GGF, manufactured in-house), and maintained at 37°C / 10% CO_2_. For stimulation and/or differentiation experiments, cells were maintained in serum-free SATO medium ^61^, consisting of low glucose DMEM (Gibco) supplemented with 100µg/ml bovine serum albumin (Gibco), 60ng/ml progesterone (Sigma), 16μg/ml putrescine (Sigma), 40ng/ml sodium selenite (Sigma), 62.5ng/ml thyroxine (Sigma), 62.5ng/ml triiodothyronine (Sigma), 20μg/ml insulin (Gibco), and 100μg/ml transferrin (R&D).

#### Proliferation assays

To assay SC proliferation *in vitro*, 2.5 x 10^4^ cells were seeded in SC medium onto coverslips coated with 20µg/ml laminin (Sigma) and 10µg/ml fibronectin (Sigma). The following day, cells were washed 3 times in SATO medium and incubated for 30h in SATO medium without insulin, containing 10µM EdU (Invitrogen), in the presence or absence of 100ng/ml NRG1 (R&D), 50ng/ml IGF1 (BioScience), and 10ng/ml TGF-β (R&D), alone or in combination. Cells were fixed in 4% paraformaldehyde/PBS for 15 mins at RT and labelled for EdU and Hoechst following the manufacturer’s instructions (Invitrogen). At least 8 different fields containing at least 1000 cells in total were counted from 2 duplicate coverslips per experiment.

#### Short interfering RNA knockdowns

Short interfering RNAs (siRNA) were purchased from Qiagen, with All-star scrambled (scr) control siRNA (Qiagen) used as a control. In brief, 2.2 x 10^5^ SCs were seeded onto PLL-coated 60mm plates. The following day, siRNAs were mixed in DMEM (Gibco) with HiPerfect transfection reagent (Qiagen) and incubated for 10 mins at RT to allow complexes to form. siRNA complexes were added to cells for 16h and washed with SC medium. For Nf1-knockdown experiments, 2 different Nf1-targeted siRNAs (Qiagen) were combined to a final concentration of 2nM, and knockdown was assessed by Western blot at 48h and 72h post-transfection.

#### Target sequences

*Nf1* siRNA 1 (*rattus norvegicus)*: 5’-CAAGCTAGAAGTGGCCTTGTA *Nf1* siRNA 2 (*rattus norvegicus)*: 5’-TGGCCTAAGATTGACGCTGTA Scrambled (scr): 5’-AATTCTCCGAACGTGTCACGT

#### Differentiation Assays

Schwann cells were grown to full confluency in SC medium, washed 3 times in SATO medium and subsequently cultured in SATO medium. 1mM dibutyryl-cAMP (dbcAMP, Sigma) was then added to induce SC differentiation in the presence or absence of 10ng/ml TGF-β (R&D) and harvested 30h after dbcAMP addition, using Trizol reagent (Invitrogen) for RNA analysis. For cells undergoing inhibitor treatments, cells were pre-treated with 0.5μM PD184352 (MedChemExpress), 10μM Galunisertib (MedChemExpress), or solvent (DMSO, Tocris) for 15 mins at 37°C / 10% CO_2_ prior to dbcAMP addition. The Δ-RafER construct in NSΔRafER SCs was activated using 100nM 4-hydroxytamoxifen (Sigma) in ethanol ^21^, added concomitantly with dbcAMP as a control for the effectiveness of MEK inhibition by PD184352 (MedChemExpress).

#### Adult mouse DRG explant cultures

Dorsal root ganglia (DRG) were dissected from 8-week old control mice for SC/axon association assays, or 8-week old Nf1^fl/fl^ or P0-Cre:Nf1^fl/fl^ mice for myelination assays. Dissected DRGs were decapsulated using tweezers, washed in Hanks’ Balanced Salt Solution (HBSS, Gibco), explanted onto coverslips pre-coated with 100μg/ml PLL (Sigma) and 50µg/ml laminin (Sigma), and incubated overnight at 37°C / 5% CO_2_ in DRG explant medium consisting of Neurobasal medium (Gibco) supplemented with 1X B27 supplement (Gibco), 100U/ml penicillin and 100µg/ml streptomycin (Gibco), 2mM L-glutamine (Gibco), 4mg/ml D-glucose (Corning), and 100ng/ml nerve growth factor (NGF, Alomone Labs). Culture medium was replenished the following day and every 2 days thereafter. For SC/axon association and dissociation assays, 1µM cytosine arabinoside (AraC, ThermoFisher) was added to the first 2 medium changes to eliminate proliferating cells.

#### SC/axon association assays

To assay the formation of SC/axonal interactions, 3 x 10^4^ SCs were seeded onto DRG coverslips on Day 6 post-explantation. For SCs that had undergone siRNA transfection, SCs were transfected 48h before seeding. Co-cultures were then incubated for 12h at 37°C / 10% CO_2_ in DRG explant medium in the presence or absence of 10ng/ml TGF-β (in 0.3% BSA 4mM HCl, R&D), 3µM SIS_3_ (in DMSO, MedChemExpress), or 0.5µM PD184352 (in DMSO, MedChemExpress). Co-cultures were then fixed in 4% PFA/PBS for 25 mins at RT, followed by immunostaining. SC/axonal interactions in DRG co-cultures were scored as previous studies ^46^.

#### SC/axon dissociation assays

To assay disruption to pre-existing SC/axonal interactions, 3 x 10^4^ SCs were seeded onto DRG coverslips on Day 6 post-explant. Co-cultures were then incubated for 48h at 37°C / 10% CO_2_ in DRG explant medium. After this time, the medium was replenished and supplemented with 10ng/ml TGF-β (R&D) or solvent control (0.3% BSA, 4mM HCl). Cultures were maintained for another 48h at 37°C / 10% CO_2_ and then fixed in 4% PFA/PBS for 25 mins at RT. SC/axonal association was scored as above.

#### Myelination assays

Isolated DRGs were maintained in DRG explant medium for 7 days at 37°C/ 5% CO_2_. DRGs were then incubated in DRG differentiation medium consisting of DMEM/F-12 medium (Gibco) supplemented with 1X N-2 supplement (Gibco), 100U/ml penicillin and 100µg/ml streptomycin (Gibco), 1X GlutaMAX supplement (Gibco), and 50ng/ml nerve growth factor (NGF, Alomone Labs) for 7 days in 37°C / 5% CO_2_, with the medium replenished every 2 days. DRGs were then incubated in DRG myelination medium consisting of Minimum Essential Medium (MEM, Gibco) supplemented with 1X N-2 supplement (Gibco), 100U/ml penicillin and 100µg/ml streptomycin (Gibco), 1X GlutaMAX supplement (Gibco), 50ng/ml nerve growth factor (NGF, Alomone Labs), 4mg/ml D-glucose (Corning), 5% horse serum (Gibco), 0.5µM forskolin (Abcam), 50µg/ml bovine pituitary extract (Gibco), 50µg/ml L-ascorbic acid (Sigma), in the presence or absence of 0.1, 1, or 10ng/ml TGF-β (in 0.3% BSA, 4 mM HCl, R&D). DRG cultures were maintained in DRG myelination medium for 21 days at 37°C / 5% CO_2_, with the medium replenished every 2 days. Cultures were then fixed in 4% PFA/PBS for 25 mins at RT, followed by immunostaining. Myelination of DRG axons by SCs was quantified using myelin basic protein (MBP) as a marker for myelin sheaths from confocal images in ImageJ/Fiji. Briefly, the extent of myelination was determined by measuring the total length of myelinated (MBP+) fibres in a field of view. At least 6 fields of view were quantified for duplicate coverslips per experiment. Each individual experiment used DRGs harvested from an individual animal.

#### Immunofluorescence (tissue)

For unfixed tissue, sciatic nerves were dissected and immediately placed into optimal cutting temperature (OCT) solution and snap-frozen in liquid nitrogen. For fixed tissue, nerves were dissected and fixed in 4% PFA/PBS overnight at 4°C, then incubated in 30% sucrose/PBS overnight at 4°C, incubated for 2h at RT in a 1:1 mix of 30% sucrose and OCT, embedded into OCT, and snap-frozen in liquid nitrogen.

Longitudinal and transverse cryo-sections (10–12µm) were obtained using a cryostat (Leica). For fixed tissue, cryo-sections were permeabilised in 0.3% Triton X-100/PBS for 30 mins at RT, blocked in 10% donkey or goat serum in PBS for at least 1h at RT, and incubated in appropriately diluted primary antibody in blocking solution overnight at 4°C. Sections were then washed 3 x 15 mins in PBS and incubated in AlexaFluor-secondary antibody (1:400) and Hoechst (1:1000) in blocking solution for 1h at RT. Sections were then washed 3 x 15 mins in PBS, rinsed in distilled H_2_O, and mounted using Fluoromount-G (Invitrogen).

For EdU labelling of tissue sections, the above protocol was followed with an additional EdU labelling step using the Click-iT EdU detection kit (Invitrogen) according to the manufacturer’s instructions. For MBP staining, 10µm transverse cryo-sections of fixed tissue were instead permeabilised in 100% Methanol (Sigma) for 25 mins at -20°C, before following the protocol above. For phospho-Smad3 staining using unfixed frozen tissue, longitudinal sections (10µm) were cut, and immediately fixed in 4% PFA /TBS for 10 mins. Sections were washed in 1mM Na_3_VO_4_ / 1mM NaF / 0.1% Triton X-100 in TBS (TBSTi) then incubated in 0.27% NH_4_Cl / 0.37% Glycine in TBSTi for 10 mins at RT. Sections were then permeabilised in 1% IGEPAL / 0.1mM tetramisol in TBSTi for 10 mins at RT. Sections were then incubated in 3% H_2_O_2_ for 30 mins at RT, blocked in 3% BSA in TBSTi for 1h at RT, and incubated in primary antibody diluted in 3% BSA in TBSTi overnight at 4°C. Sections were then washed 3 times in TBSTi and incubated in HRP-conjugated secondary antibody for 1h at RT. Phospho-smad3 (Abcam) was detected using a Tyramide amplification kit (Invitrogen) according to the manufacturer’s instructions. After phospho-Smad3 detection, sections were incubated in AlexaFluor-conjugated secondary antibody (1:400) and Hoechst (1:1000) in 3% BSA in TBS for 1h at RT. Sections were then washed 3 times in TBS, rinsed in distilled H_2_O, and mounted using Fluoromount-G (Invitrogen).

#### Immunofluorescence (cells)

SC cultures were fixed in 4% PFA/PBS for 15 mins at RT, and DRG cultures were fixed in 4% PFA/PBS for 25 mins at RT. In both cases, coverslips were permeabilised in 0.3% Triton X-100/PBS for 20 mins at RT, and blocked in 3% BSA/PBS for at least 1h at RT. Coverslips were then incubated in appropriately diluted primary antibodies in 3% BSA overnight at 4°C. Coverslips were then washed in PBS, and incubated in secondary antibodies (1:400), Hoechst (1:1000), and phalloidin-546 (1:1000) in 3% BSA for 1h at RT. For proliferation assays, EdU was detected using the Click-iT EdU detection kit (Invitrogen). Coverslips were then washed in PBS, rinsed in distilled H_2_O, and mounted using Fluoromount-G (Invitrogen).

#### Immunofluorescence antibodies

Antibodies used for immunofluorescence at indicated dilutions: p75NTR (1:300, Sigma-Aldrich), GFP (1:1000, Abcam), S100 (1:100, DAKO), tdTomato (1:200, OriGene), phosphorylated Smad3 (1:1000, Abcam), Glut1 (1:1000, Abcam), Neurofilament (1:1000, Abcam), CD31/PECAM-1 (1:100, R&D), F4/80 (1:400, Bio-Rad), Iba1 (1:500, WAKO), CD117 (1:100, R&D), CD3 (1:200, Invitrogen), CD45R (1:200, Invitrogen), Ly6G (1:200, BioLegend), MBP (1:2000, ThemoFisher). AlexaFluor-conjugated secondary antibodies were purchased from ThermoFisher or Abcam and used at a dilution of 1:400.

#### Hybridisation chain reaction (HCR)

For *in situ* HCR, longitudinal cryo-sections (10-12µm) of fixed, frozen sciatic nerve were obtained using a cryostat (Leica). Sections were washed twice in diethyl pyrocarbonate (DEPC)-treated PBS for 5 mins and permeabilised in 70% Ethanol in DEPC-treated H_2_O for 3h at 4°C. For hybridisation, sections were incubated in Hybridisation Buffer (Molecular Instruments) for 30 mins at 37°C, before incubating with RNA Probe Solution (Molecular Instruments) targeting either *tdTomato* or *Tgfb1* mRNA for 16h at 37°C. Samples were then washed 5 times in Probe Wash Buffer (Molecular Instruments) for 15 mins. For amplification, sections were incubated in Amplification Buffer (Molecular Instruments) for 30 mins at RT. Hairpin solution was prepared by snap-heating hairpins at 95°C for 90 sec, allowed to cool to RT for 30 mins in the dark, and diluted 1:200 in Amplification Buffer (Molecular Instruments). Sections were incubated in hairpin solution for 16h at RT in the dark. Sections were then washed 4 times in 5X SSCT solution. To co-immunostain, sections were post-fixed in 4% PFA/PBS for 10 mins at RT. Samples were then washed twice with DEPC-PBS, blocked in 5% donkey serum for 1h at RT, and incubated in primary antibody diluted in blocking solution overnight at 4°C. Sections were then washed 3 times in DEPC-PBS, incubated in AlexaFluor-conjugated secondary antibody diluted in blocking solution for 1h at RT. Subsequently, samples were washed 3 times in DEPC/PBS, incubated in DAPI (1:500 in DEPC-PBS) for 30 mins at RT, washed twice in DEPC/PBS, and mounted using Fluoromount-G (Invitrogen).

#### Confocal microscopy

Confocal microscopy was performed using Leica SPE3, Leica TCS SP8 STED, and Zeiss LSM880 Multiphoton microscopes. Leica SPE3 was used with the 63X (1.3 NA) and 40X (1.15 NA) oil immersion objectives. Leica TCS SP8 STED and Zeiss LSM880 Multiphoton were used with the 20X (0.8 NA) objective. The Z-step size in all images ranged between 0.2 and 1µm. In cases using tile scan images, individual tiles were stitched automatically using LAS X software (Leica).

#### Image analysis

Confocal images were opened and processed using ImageJ/Fiji and Photoshop Software (Adobe). Parameters for optimisation of visualisation were applied to all images within an individual experiment.

#### Proliferation

To determine the proliferation rates of cell types of interest, EdU labelling was performed on cryo-sections of EdU-injected mice. For all images, a z-projection of 10µm with 0.5µm increments was used, and the number of Hoechst+ and Hoechst+/EdU+ cells in each field was counted. Proliferation was expressed as the percentage of EdU+ cells. At least 4 fields (∼40 cells/field) of each region were counted per animal.

#### pSmad3 fluorescence intensity

To determine the fluorescence intensity of pSmad3 staining per cell or per SC, a 10µm stack was imaged with 0.2µm increments. The pSmad3 signal was thresholded to background and the pSmad3 intensity of each cell measured by creating a Hoechst mask. For recombined SCs, the tdTomato channel was overlayed, and all tdTomato negative nuclei were omitted from the quantification. At least 4 fields (∼40 cells/field) of each region of interest per animal was analysed.

#### Nerve width and tumour area

To measure tumour size, nerves harvested at 6-months post-injury were positioned under a Leica DFC420C system so that the uninjured and injured fascicles could be imaged side by side. The maximum nerve width (transverse) of the injury site was measured on ImageJ/Fiji.

#### Tumour area

The tumour area was measured by two methods: from macroscopic images obtained on a Leica DFC420C system and from confocal tile-scan IF images obtained on a Leica TCS SP8 STED microscope at 20X magnification. In both cases area was measured using ImageJ/Fiji. From macroscopic images, the tumour region was considered as tissue which protruded outwards from the injury site of the sciatic nerve. Adipose and muscle tissue was visually discernible and not considered in any quantification. From IF images, the tumour region was considered as any tissue which protruded from the injury site and any tdTomato+ tissue outside of the perineurium. Both males and females were used in these quantifications, as no difference in sciatic nerve thickness could be detected between sexes.

#### SC/Axon association

SC/axon association at 6 months post-injury was quantified from transverse IF images as the percentage of recombined (tdTomato+) SCs that displayed evident contact with an axon based on proximity and morphology, using Neurofilament (NF) as an axonal marker. A 10µm stack was imaged (0.2µm increments) for each field. At least 4 fields of each region were counted per animal, with each field approximately 60 – 100 tdTomato+ cells.

#### SC myelination

SC myelination at 6 months post-injury was quantified from transverse IF images as the percentage of recombined (tdTomato+) SCs that displayed evident myelinating SC morphology and were positive for the myelin marker Myelin Basic Protein (MBP). A 10µm stack was imaged (0.2µm increments), and cells were only considered negative for MBP if no myelin was visible at any point in the stack. At least 4 fields of each region were counted per animal, with each field approximately 60 – 100 tdTomato+ cells.

#### Enclosed SC proportion

The proportion of SCs enclosed by Glut1+ perineurial cells at 6 months post-injury was quantified from transverse IF images as the percentage of recombined (tdTomato+) SCs residing within a Glut1+ minifascicle in each field. A 10µm stack was imaged (0.2µm increments), and cells were only considered unenclosed if they remained outside of a Glut1+ minifascicle over the entire stack. At least 4 fields of each region were counted per animal, with each field approximately 60 – 100 tdTomato+ cells.

#### Axon and BV coverage

Blood vessel / axonal coverage was quantified from confocal tile scan images stained with CD31 (blood vessels) or Neurofilament (axons). The channel of interest was thresholded to background and converted to a binary mask. The area of the selection within a region was then used to measure blood vessel or axonal coverage. The entire tile scan was measured per animal.

#### Axon deviation / BV deviation

The deviation of blood vessels and axons from their normal migratory path was quantified in tile scan images of longitudinal sections of nerves using ImageJ/Fiji. A z-projection of 10µm with 0.5µm increments was used. The region of interest was aligned vertically to the proximal-to-distal axis, and each blood vessel/axon was traced from the proximal-to-distal direction. For each blood vessel/axon, the angle was recorded, and the deviation measured either side of the proximal-to-distal axis. At least 3 fields of each region were counted per animal.

#### Cell density measurements

Cell density was quantified from confocal microscopy images using Fiji. For each image, a z-projection of 10µm with 0.5µm increments was used, and the number of Hoechst+ cells positive for a specific marker (macrophages – F4/80 or Iba1; mast cells – CD117; T cells – CD3; neutrophils – Ly6G; perineurial cells – Glut1) in each field was counted. Cell density was expressed either as number of cells per field or number of cells per mm^2^. At least 4 fields of each region (∼40 cells) were counted per animal.

#### Escaped region quantifications

The escaped region area at 14 days post-injury was quantified using Fiji from confocal tile scan images of longitudinal sections and was defined as the region of regenerated tissue that was not enclosed by Glut1+ cells. Adipose and muscle tissue were excluded from the quantification. The entire region of interest in the tile scan was measured per animal. Escaped SC number was measured using the same images.

#### FiSH coverage

The location of the *Tgfb1* mRNA signal at 14 days post-injury was quantified from confocal tile scan images probed for *Tgfb1* mRNA and immunostained for Glut1 (perineurial cells). Glut1 staining was used to define the regenerating and escaped region. The *Tgfb1* channel of interest was thresholded to background and round objects selected using the Fiji selection tool corresponding to punctate staining. A minimum size threshold for objects was used to improve object recognition. Autofluorescent erythrocytes were excluded from the quantification. The coverage of *Tgfb1* mRNA was expressed as density of puncta within each region of interest. At least 6 fields of each region were counted per animal.

#### FiSH puncta per cell

To measure the number of *Tgfb1* puncta per cell, the puncta within each cell type was counted using 0.2µm increments over 12µm z-stacks. Only cells for which the entire cytoplasm and nucleus could be identified within the 12µm stack were considered. Relevant cell markers were used to label each cell type (macrophages – Iba1; recombined SCs – tdTomato). At least 5 fields of each region were counted per animal, with at least 30 cells counted per animal.

#### DRG axon/SC interactions – ex vivo

Schwann cell/axonal interactions in DRG co-cultures were scored as previously described ^46^. Cultures were imaged at high magnification using confocal microscopy, and SCs were individually scored based on their alignment to nearby axons. SCs that displayed tight cytoplasmic and nuclear alignment to a contacting axon were considered ‘associated and aligned’; SCs that failed to contact an axon or failed to align their nucleus and cytoplasm to an axon were not considered to be correctly associated. At least 400 cells were counted across multiple fields from at least 2 duplicate coverslips for each condition in each independent experiment. Fields used for quantification were acquired at least 200µm away from the DRG cell bodies.

#### DRG axon myelination – ex vivo

Myelination of DRG axons by SCs was quantified using myelin basic protein (MBP) as a marker for myelin sheaths from confocal images using Fiji. The extent of myelination was determined by measuring the total length of myelinated (MBP+) fibres within a field of view. For each experiment, the mean was calculated from at least 6 fields imaged per coverslip, using a minimum of 2 duplicate coverslips for each condition in each independent experiment.

#### RNA extraction from tissue and cultured cells

Sciatic nerves were dissected at indicated timepoints and snap-frozen in liquid nitrogen. Samples were then crushed and homogenised on dry ice and lysed using 200µl Trizol reagent (Invitrogen). A further 300µl of Trizol reagent was then added and incubated for 5 mins at RT. After lysis, 100µl of chloroform was added, the samples vortexed for 2 mins, and centrifuged for 15 mins at 4°C. The upper RNA phase was transferred to a fresh tube, and 500µl of 2-propanol was added to the sample for 10 mins at RT and centrifuged for 10 mins at 4°C to precipitate RNA. The supernatant was discarded, and the RNA pellet was washed 2 times with 80% ethanol by centrifuging for 5 mins at 4°C. After washes, the pellet was air-dried for 10 mins and resuspended in nuclease-free H_2_O. RNA was quantified by Nanodrop measurement. Cultured SCs were lysed directly in Trizol reagent. After lysis, RNA was extracted and quantified following the same protocol as above.

### RNA-seq

#### RNA extraction for RNA-seq

Sciatic nerves fragments were lysed using the same protocol as described above using Trizol reagent and phase separation using chloroform. 6 nerve fragments (each fragment 0.5mm) were combined from separate mice for each sample. Only mice which exhibited a recombination rate of at least 30% were used. 3 independent samples were obtained for each region and genotype of interest. After phase separation, the RNA phase was transferred to a fresh tube, and 70% ethanol added in a 1:1 mix (final ethanol concentration of 35%). RNA was then extracted using PureLink RNA Micro Scale kit columns (Invitrogen, 12183016) according to the manufacturer’s instructions. An additional DNA removal step was performed to eliminate genomic DNA using a RNase-free DNase kit (Qiagen) according to the manufacturer’s instructions. RNA was eluted in nuclease-free H_2_O and quantified by Nanodrop measurement.

#### Ribosomal depletion, library preparation and RNA sequencing

Ribosomal depletion, library depletion and RNA sequencing was performed by the Genomic Facility at the University of Liverpool, UK. Briefly, ribosomal RNA (rRNA) was depleted with the Ribo-Zero Gold Kit (Illumina) and extent of rRNA depletion was analysed with Bioanalyzer RNA pico chip. The NEB Ultra Directional RNA Library Preparation Kit (NewEnglandBiolabs, E7420) was used to prepare the RNA-Seq libraries from the enriched material. All the enriched material was used and following 15 cycles of amplification, the RNA was purified using AMPure XP beads (Beckman Coulter, A63882) The libraries were quantified using Qubit and the size distribution was analysed using the Agilent 2100 Bioanalyser. The final libraries were pooled in equimolar amounts. The quantity and quality of the libraries were determined by Bioanalyzer and subsequently by RT-qPCR using the Illumina Library Quantification Kit from Kapa (KK4854) on a Roche Light Cycler LC480II. The template DNA was denatured and loaded at 300pM. The sequencing was carried out on two lanes of an Illumina HiSeq4000 at 2x150 bp paired-end sequencing ^62,63^.

#### RNA-seq analysis

Prior to the bioinformatics analysis, the quality of the raw data was assessed, and the data was normalised. The initial raw data processing steps (basecalling, de-multiplexing and trimming) were performed at the Genomic Facility of the University of Liverpool. Basecalling and de-multiplexing of the indexed reads was performed using CASAVA version 1.8.2. The raw fastq files were trimmed using Cutadapt version 1.2.1. Low quality bases were removed from the reads by using Sickle version 1.200 with a minimum window quality score of 20. After trimming was completed, reads shorter than 10bp were removed. Paired-end reads were mapped to the Ensembl mouse transcriptome reference sequence (mm10). Mapping and generation of read counts per transcript were performed using Kallisto, based on pseudoalignment. R/Bioconductor package Tximport was used to import the mapped counts data and summarise the transcripts-level data into gene-level data. DESeq2 (Anders and Huber, 2010; Love et al., 2014) and SARTools packages (developed at PF2, Institute Pasteur) were used for differential gene expression analysis, using a false discovery rate (FDR) <0.05, following normalisation with a negative binomial generalised linear model.

#### Reverse transcription and qPCR

For reverse transcription, 1µg of RNA was used to synthesise complementary DNA using the Superscript II Reverse Transcriptase kit (Invitrogen). For qPCR, the SYBR Select Master Mix kit was used (ThermoFisher). A total of 40 cycles were run in the following steps: 95°C for 20 sec, 58°C for 30 sec, 72°C for 20 sec. Relative expression values for each gene of interest were obtained by normalising to B2M and expressed as a fold change relative to the control condition. The following primer sequences were used:

*B2m (rattus norvegicus)*

Fwd: 5’- TGACCGTGATCTTTCTGGTG

Rev: 5’- ATTTGAGGTGGGTGGAACTG

*Krox20/Egr2 (rattus norvegicus)*

Fwd: 5’- CAGTACCCTGGTGCCAGCTG

Rev: 5’- TGTGGATCTCTCTGGCACGG

*Mbp (rattus norvegicus)*

Fwd: 5’- CACAAGAACTACCCACTACGG

Rev: 5’- GGGTGTACGAGGTGTCACAA

*Prx (rattus norvegicus)*

Fwd: 5’- GGAACTCTGGAGGTGTCTGG

Rev: 5’- TCCTGGGTGATGGTCTTTTC

*B2m (mus musculus)*

Fwd: 5’- CAGTCTCAGTGGGGTGAAT

Rev: 5’- ATGGGAAGCCGAACATACTG

*Cxcl14 (mus musculus)*

Fwd: 5’- GTTATCGTCACCACCAAG

Rev: 5’- CTCTCAACTGGCCTGGAGT

*Cilp (mus musculus)*

Fwd: 5’- GTTCCGAGTTCCTGGCTTGT

Rev: 5’- AATGTATGGGGTCTCTGCCC

*Vcan (mus musculus)*

Fwd: 5’- TCCTGATTGGCATTAGTGAAG

Rev: 5’- CTGGTCTCCGCTGTATCC

*Tnfsf18 (mus musculus)*

Fwd: 5’- TCAAGTCCTCAAAGGGCAGAG

Rev: 5’- GGCAGTTGGCTTGAGTGAAGTA

*IL6 (mus musculus)*

Fwd: 5’- GAGGATACCACTCCCAACAGACC

Rev: 5’- AAGTGCATCATCGTTGTTCATACA

*Edil3 (mus musculus)*

Fwd: 5’- CTGTGAGTGTCCAGAAGGCT

Rev: 5’- AGGCTTCGCTTATCTCACAGG

*Slitrk6 (mus musculus)*

Fwd: 5’- GCTTTACAGAGTCAGATCCAATCAT

Rev: 5’- AAAGAGTGTCGCAAGAGCCT

*Tgfb2 (mus musculus)*

Fwd: 5’- CTCCCCTCCGAAACTGTCTG

Rev: 5’- TGTCTGGAGCAAAAGCTG

#### Western blotting

For protein analysis, SCs were washed in ice-cold PBS and lysed using a RIPA buffer consisting of 1% Triton X-100, 0.5% sodium deoxycholate, 50mM Tris pH 7.5, 100mM NaCl, 1mM EGTA pH 8, 20mM NaF, 100µg/ml PMSF, 15µg/ml aprotonin, 1mM Na_3_VO_4_, and 1:100 protease and phosphatase inhibitor cocktails (Sigma). Cells were lysed on ice for 30 mins, during which they were briefly vortexed every 10 mins, and homogenised using a 26-gauge needle. Lysates were centrifuged at 750 x g for 15 mins at 4°C, the supernatant was collected and quantified by BCA assay (Pierce, ThermoFisher), appropriately diluted in Laemmli buffer, and boiled at 95°C for 5 mins.

Western blotting was performed using Invitrogen Western blot electrophoresis systems. Protein samples (30 – 40µg) were resolved using a sodium dodecyl sulphate polyacrylamide gel electrophoresis (SDS-PAGE). Protein was transferred onto a PVDF membrane (Millipore-Immobilon) and blocked for 1h at RT in 5% milk diluted in 0.1% Tween in TBS (TBST). The membrane was incubated with primary antibodies (1:1000) diluted in 5% milk-TBST overnight at 4°C. The following day, the membrane was washed 3 times in TBST for 15 mins each, followed by incubation with the appropriate HRP-conjugated secondary antibody (1:5000) in 5% milk-TBST for 1h at RT. Subsequently, membranes were washed 3 times with TBST before detection of proteins of interest with Pierce-ECL Western blot substrate (ThermoFisher) or Luminata Crescendo Western HRP substrate (EMD-Millipore) on an Alliance Uvitec Imaging system.

#### Western blot antibodies

Antibodies used for Western blotting at indicated dilutions: Neurofibromin (1:1000, Santa Cruz), Vinculin (1:1000, Sigma-Aldrich). HRP-conjugated secondary antibodies were purchased from Cell Signaling Technology and used at a dilution of 1:5000.

### Statistical analysis

Statistical analysis was performed using Graphpad Prism software. Unless stated otherwise, data is presented as mean ± SEM and is representative of at least 3 animals or independent experiments. Data were analysed by unpaired two-tailed t-test, one-way ANOVA, or two-way ANOVA, as appropriate, followed by multiple comparisons tests, when applicable, as indicated by the Figure Legends. In all cases, statistical significance was considered at p<0.05, and further indication of the p-value is indicated by (*): *p<0.05, **<0.01, ***<0.001, ****<0.0001, ns=not significant (p>0.05). In some cases, the recorded p-value is shown. Not all comparisons are shown. The number of animals or biological repeats used in each experiment is stated in the Figure Legends.

### Supplementary Figure Legends

**Supplementary Figure S1 – related to Figure 1.**
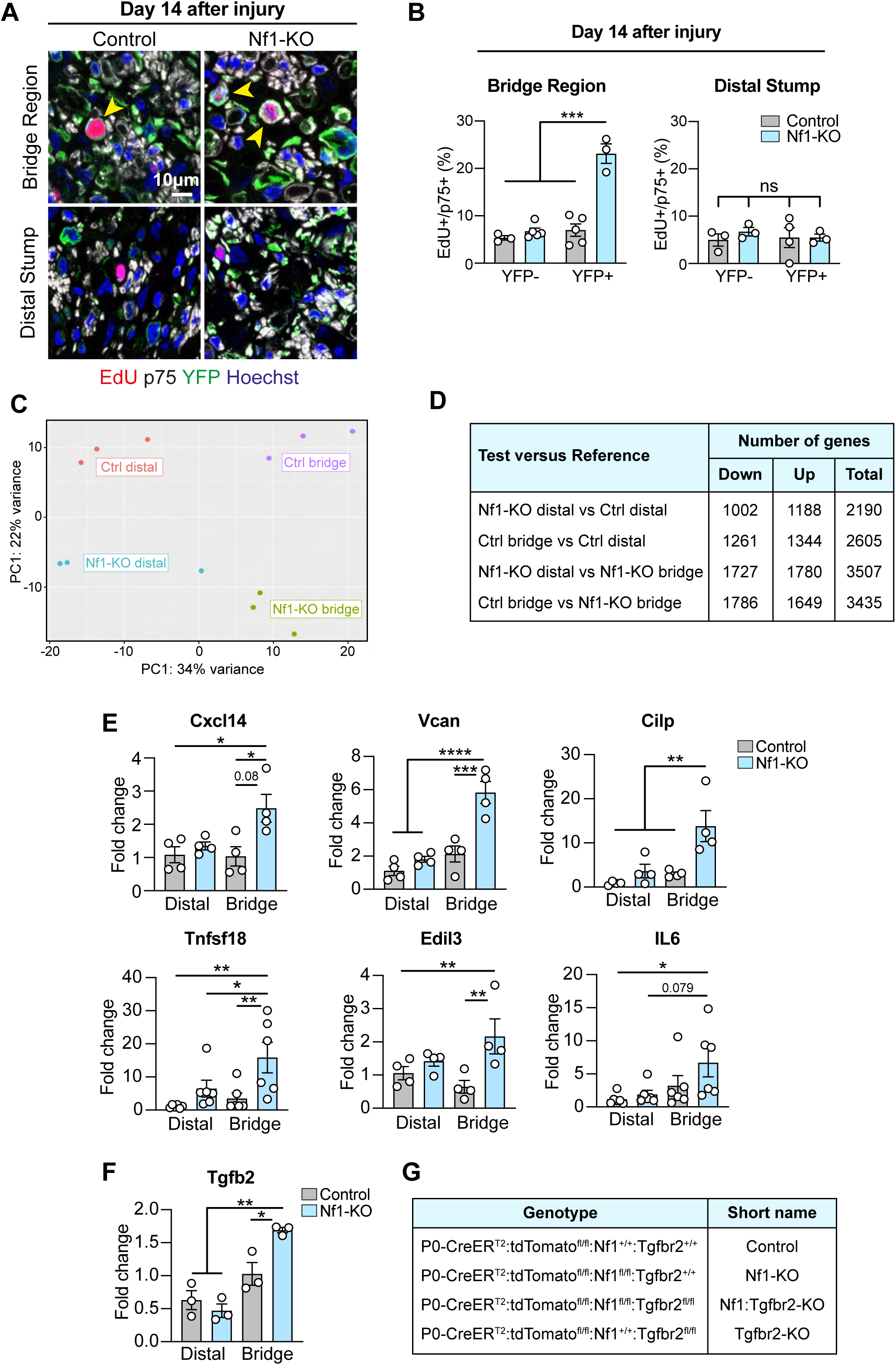
**A**. Representative confocal images of transverse sections of the bridge and distal regions from Control or Nf1-KO mice at Day 14 post-injury, showing recombined SCs (YFP, green), and labelled to detect SCs (p75, white), EdU (red), and nuclei (Hoechst, blue). Yellow arrowheads indicate EdU+ recombined SCs. **B.** Quantification of (**A**) showing the percentage of EdU+ recombined (p75+/YFP+) and unrecombined (p75+/YFP-) SCs. Each dot represents one mouse (n=3-5). Data shown as mean ± SEM. **C.** Principal Component Analysis Plot of RNA-seq samples collected from the bridge or distal regions of nerves from Control or Nf1-KO animals at Day 14 post-injury (n=3 independent RNA-seq samples, with each sample containing pooled nerve fragments from at least 6 mice). **D.** Table indicating the number of differentially regulated genes between conditions. **E.** Graphs show RT-qPCR analysis of TGF-β-targets of RNA isolated from the distal or bridge nerve regions from Control or Nf1-KO mice at Day 14 post-injury. Each dot represents one mouse (n=3-6). Data was normalised to B2m expression and shown as mean ± SEM. **F.** Graphs show RT-qPCR analysis of relative Tgfb2 mRNA levels in the distal or bridge regions of nerves from Control or Nf1-KO mice at Day 14 post-injury, showing. Each dot represents one mouse (n=3). Data is normalised to B2m and presented as mean ± SEM. **G.** Table indicating the nomenclature of the 4 different genotypes used in the study. For B, E, and F, a two-way ANOVA was used. *p<0.05, **p<0.01, ***p<0.001, ****p<0.0001, ns=not significant.

**Supplementary Figure S2 – related to Figure 2.**
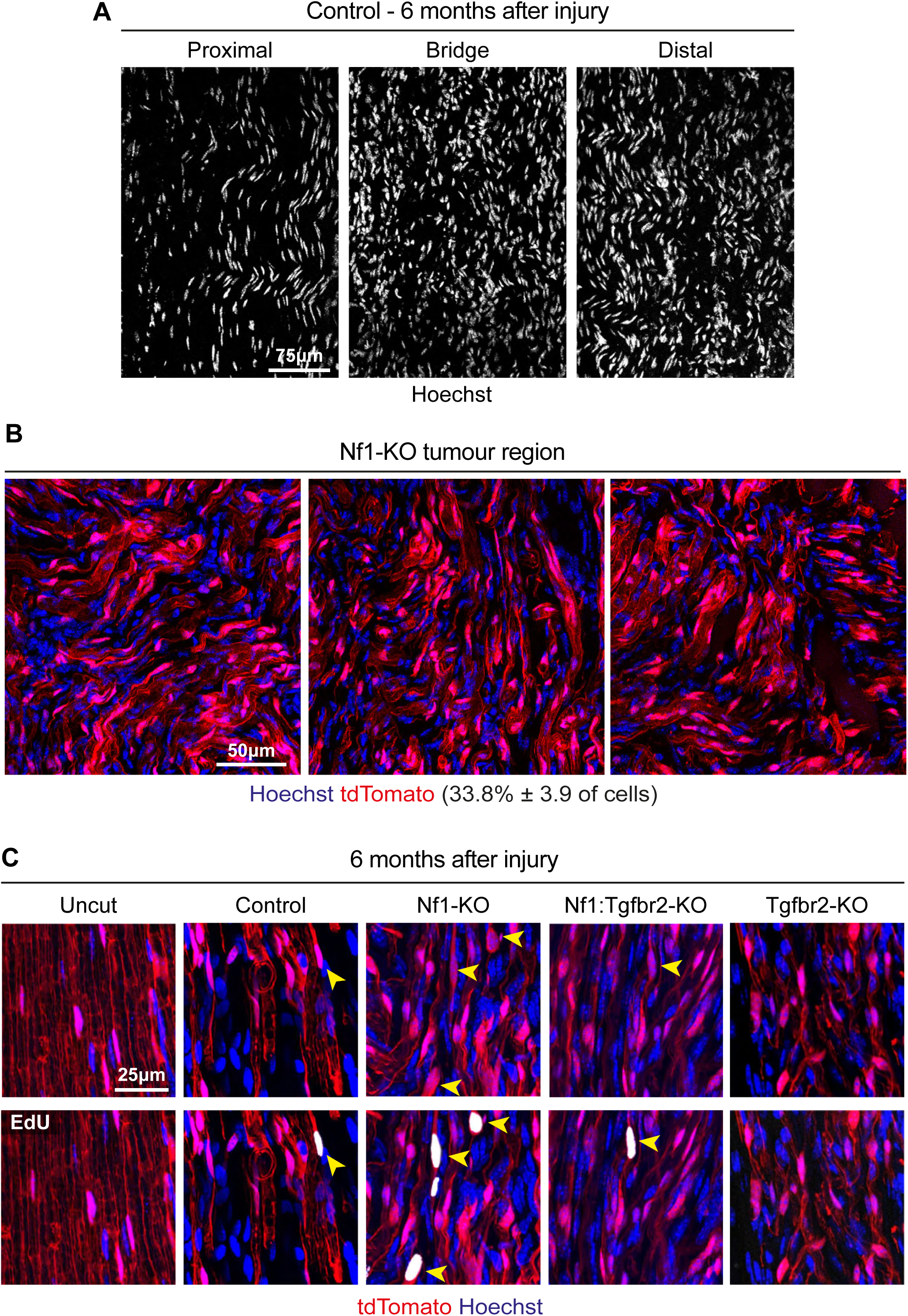
**A.** Representative confocal images of longitudinal sections of indicated nerve regions from a control mouse, 6 months after injury, labelled to detect nuclei (Hoechst, white). **B.** Representative confocal images of longitudinal sections of tumour from Nf1-KO mice, at 6 months post-injury, showing recombined SCs (tdTomato, red) and labelled to detect nuclei (Hoechst, blue). % tdTomato cells are presented as mean ± SEM (n=6 mice). **C.** Representative confocal images of longitudinal sections of the bridge region from indicated genotypes, harvested 6 months after injury, showing recombined SCs (tdTomato, red), and labelled to detect EdU (white), and nuclei (Hoechst, blue). Mice were injected with EdU 48h, 24h, and 4h before harvest. Arrowheads indicate EdU+ recombined SCs.

**Supplementary Figure S3 – related to Figure 3.**
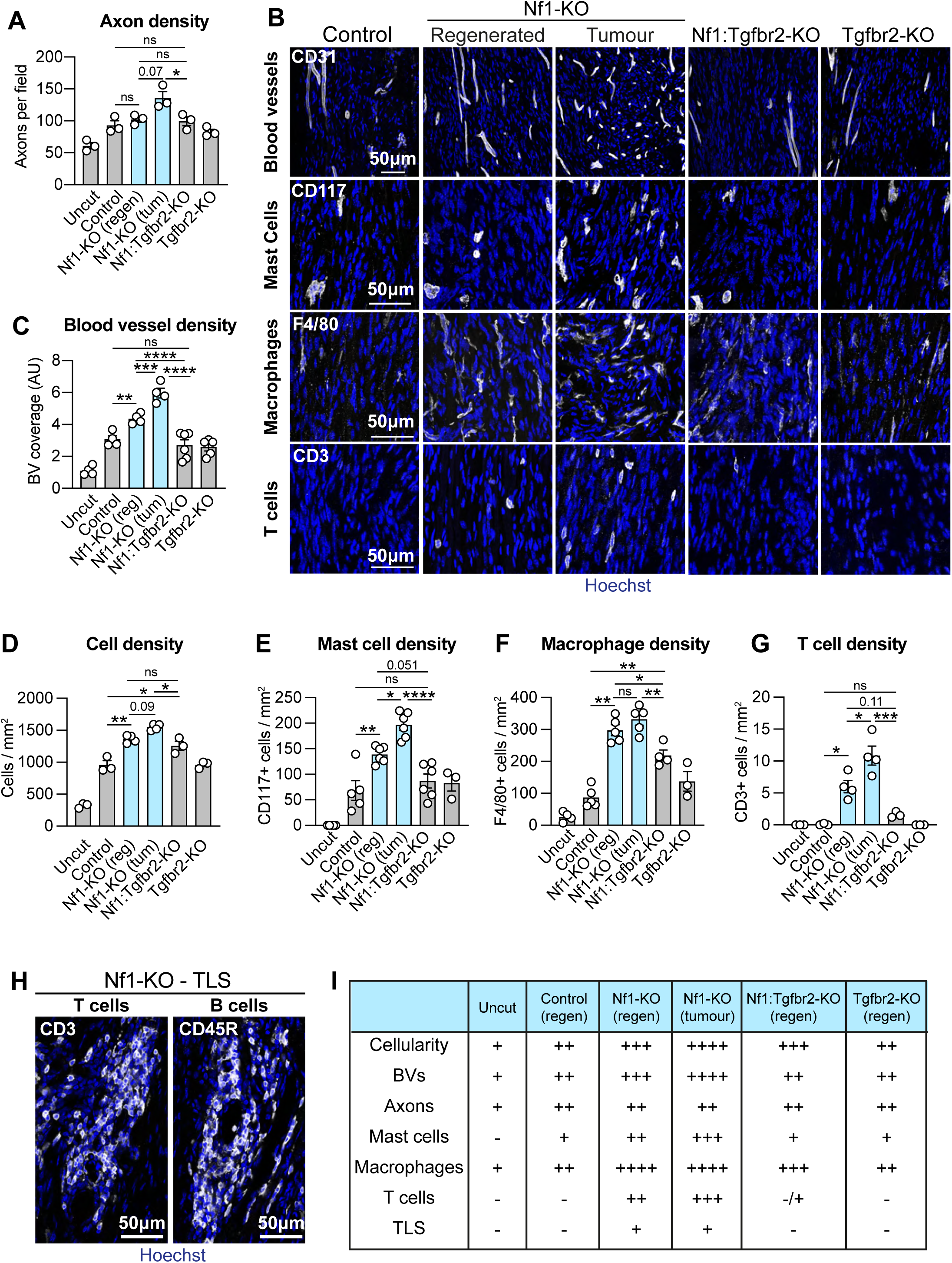
**A.** Quantification of axon density in indicated genotypes and regions, 6 months after injury. Each dot represents one mouse (n=3), data is presented as mean ± SEM **B.** Representative confocal images of longitudinal nerve sections from indicated mice at 6 months post-injury, labelled to detect blood vessels (CD31, white), mast cells (CD117, white), macrophages (F4/80, white), T cells (CD3, white), and nuclei (Hoechst, blue). Quantification of (**B**) showing (**C)** the density of blood vessels, (**D**) total cells, (**E**) mast cells, (**F**) macrophages, and (**G**) T cells, in indicated genotypes, 6 months after injury. Each dot represents one mouse. Data is presented as mean ± SEM. **H.** Representative confocal images of tertiary lymphoid structures in nerves from Nf1-KO mice at 6 months post-injury, labelled to detect T cells (CD3, white), B cells (CD45R, white), and nuclei (Hoechst, blue). **I.** Summary table of the differences between genotypes (and regions) at 6 months post-injury. For A and C-G, a two-way ANOVA was used. *p<0.05, **p<0.01, ***p<0.001, ****p<0.0001, ns=not significant.

**Supplementary Figure S4 – related to Figure 4.**
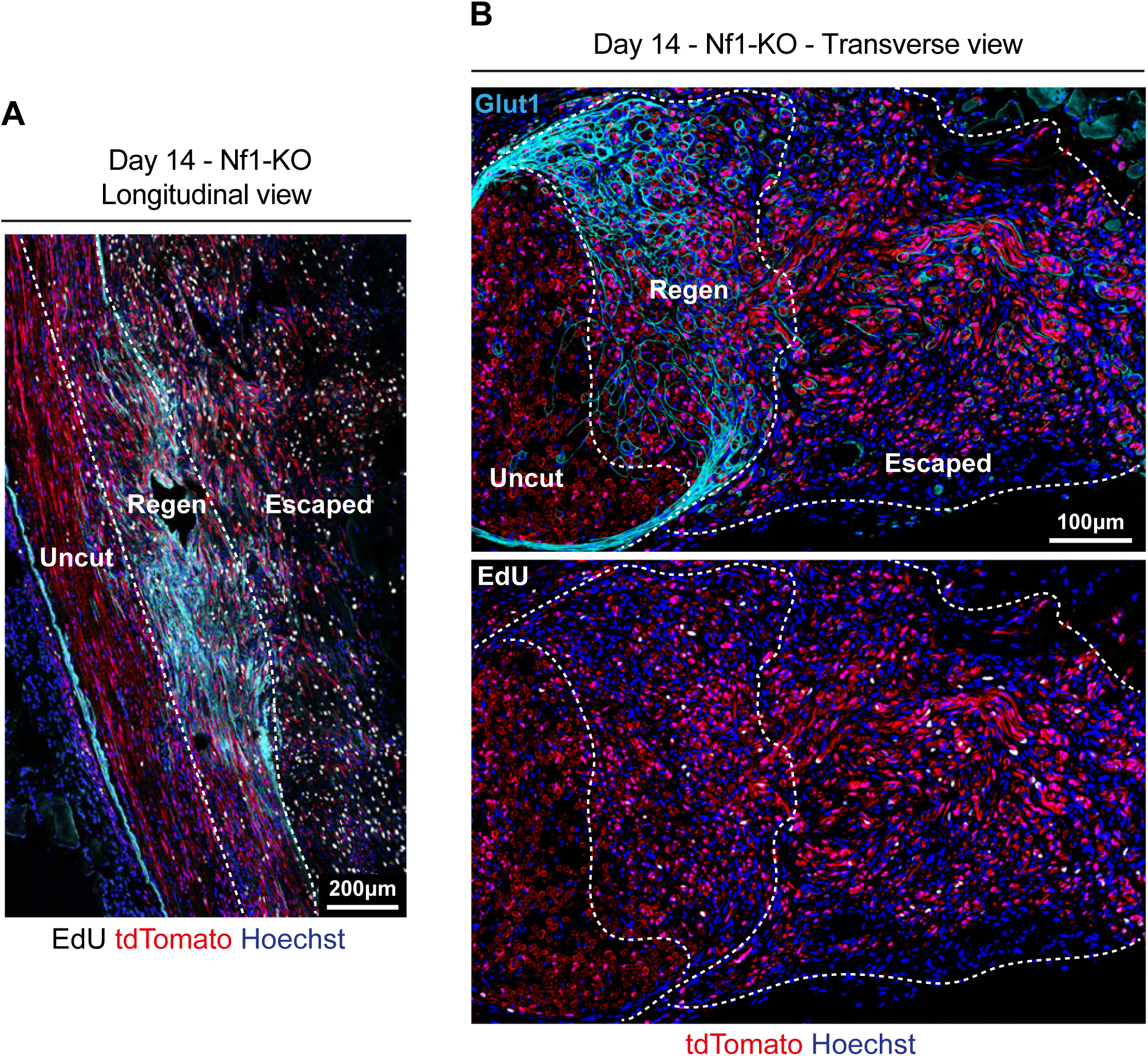
Representative tile scan images of longitudinal (**A**) and transverse (**B**) sections of nerves from Nf1-KO mice, harvested at Day 14 post-injury. EdU was injected 6h and 3h prior to harvesting. Sections show recombined SCs (tdTomato, red), and are labelled to detect EdU (white), perineurial cells (Glut1, cyan), and nuclei (Hoechst, blue). Dashed lines delineate indicated regions.

**Supplementary Figure S5 – related to Figure 5.**
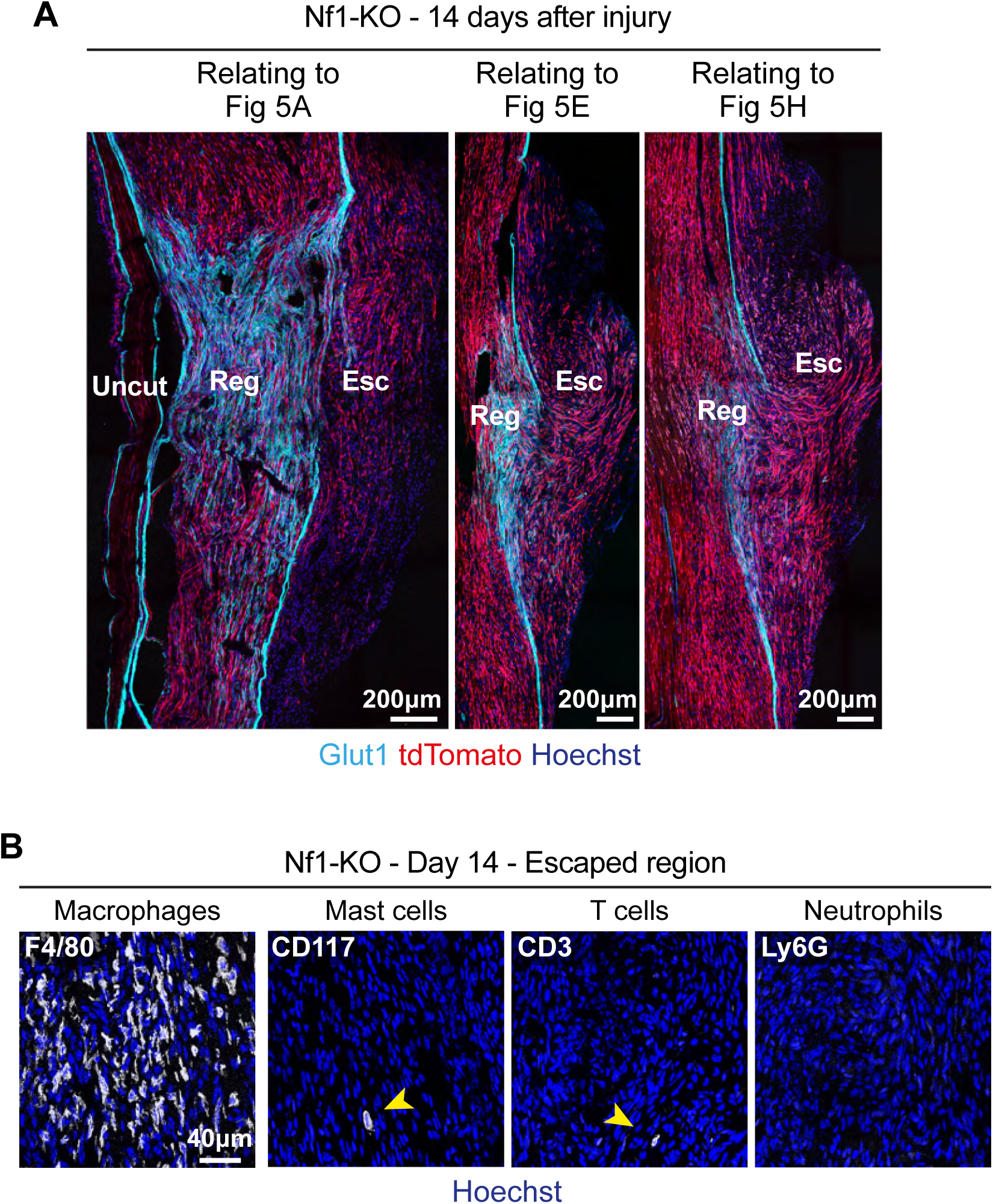
**A.** Representative tile scan confocal images of longitudinal sections of regenerated and escaped regions of nerves from Nf1-KO mice at Day 14 post-injury, showing recombined SCs (tdTomato, red), and labelled to detect perineurial cells (Glut1, blue), and nuclei (Hoechst, blue). **B.** Representative confocal images of longitudinal sections of the escaped region in nerves from Nf1-KO mice at Day 14 post-injury, labelled to detect macrophages (F4/80, white), mast cells (CD117, white), T cells (CD3, white), neutrophils (Ly6G, white), and nuclei (Hoechst, blue). Arrowheads indicate rare mast cells and T cells.

**Supplementary Figure S6 – related to Figure 6.**
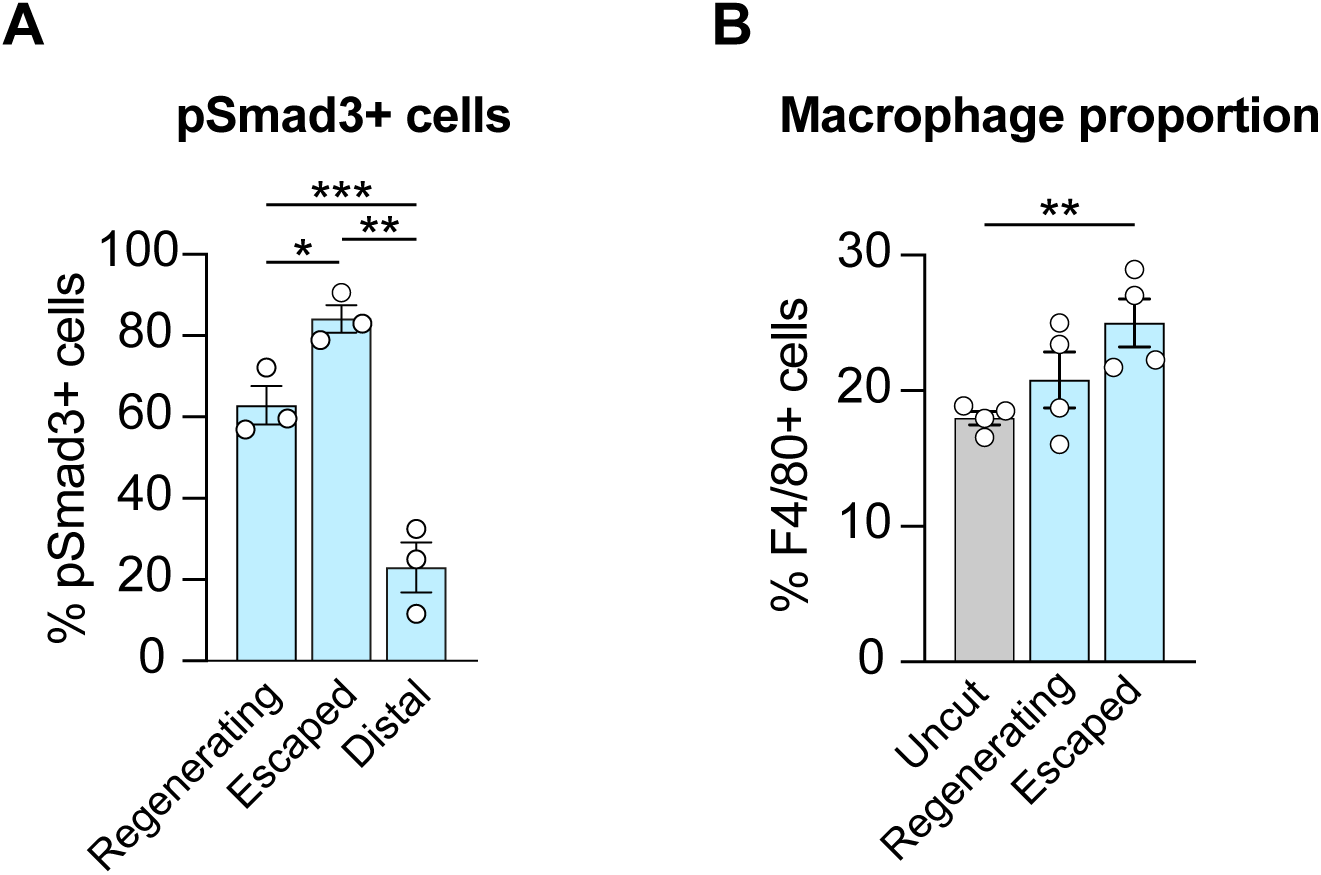
**A.** Quantification of the percentage of phosphorylated-Smad3+ cells in the indicated regions of nerves from Nf1-KO mice harvested at Day 14 post-injury. Each dot represents one mouse (n=3), data shown as mean ± SEM. **B.** Quantification of the proportion of macrophages (F4/80+) cells in the indicated regions of nerves from Nf1-KO mice, harvested at Day 14 post-injury. Each dot represents one mouse (n=4), data shown as mean ± SEM. For A and B, a one-way ANOVA was used. *p<0.05, **p<0.01, ***p<0.001.

**Supplementary Figure S7 – related to Figure 7.**
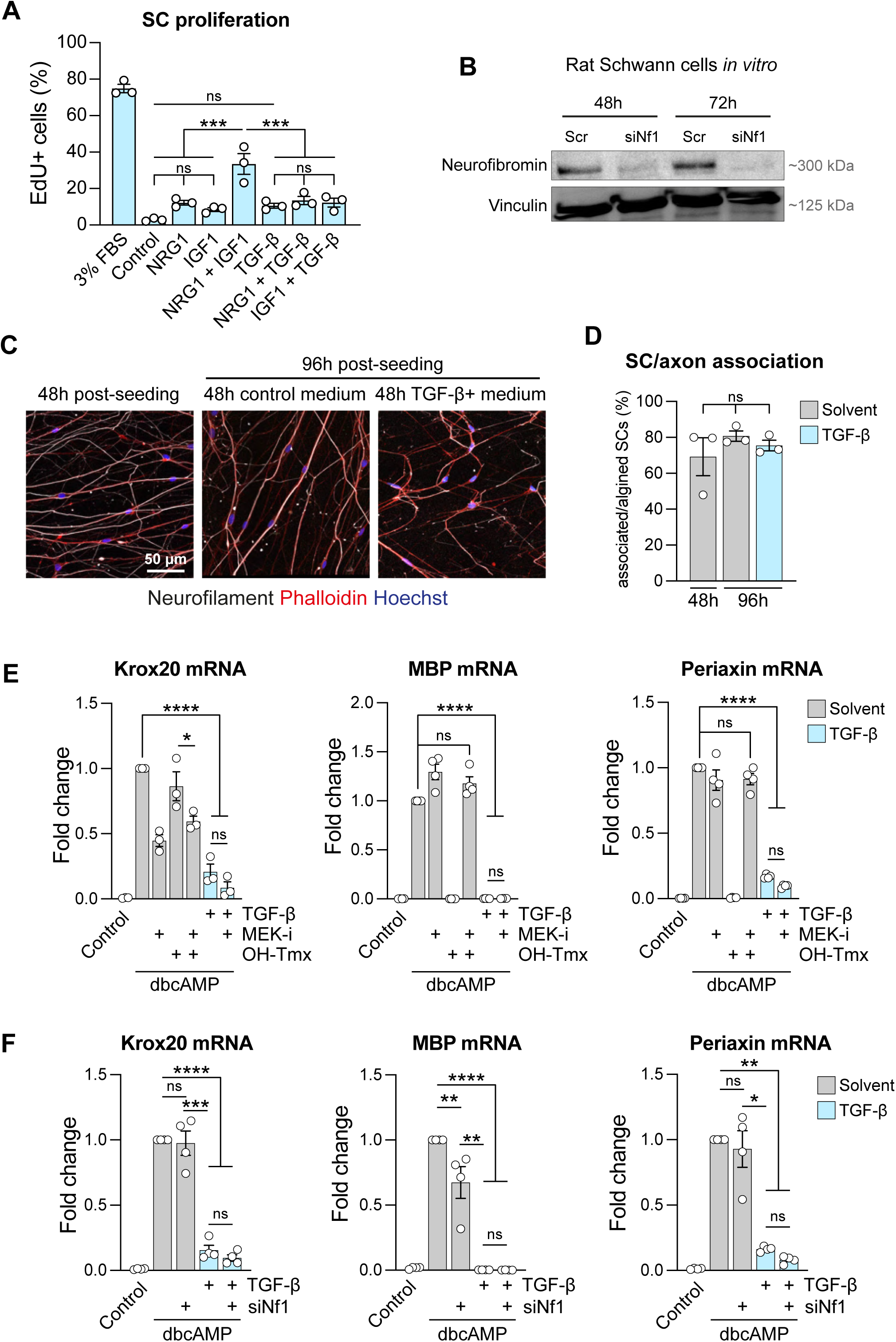
**A.** Graph shows the percentage of EdU+ cells in SCs cultured for 30h in defined serum-free medium (SATO) supplemented with NRG1 (100ng/ml), IGF-1 (50ng/ml), TGF-β (10ng/ml), alone or in combination. Serum-containing medium (3% FBS) was used as a positive control. Each dot represents one independent experiment, data shown as mean ± SEM of 3 independent experiments. **B.** Representative western blot of protein harvested from rat SCs 48h or 72h after transfection with 2nM scrambled or Nf1-targeting siRNAs. Vinculin was used as a loading control. Blot is representative of 3 independent experiments **C.** Representative confocal images of DRG/ SC co-cultures. SCs were added to DRG axons for 48h to allow robust SC/axon association, then cultured for an additional 48h in the presence or absence of 10ng/ml TGF-β. Cultures were labelled to detect axons (NF, white), actin (phalloidin-546, red), and nuclei (Hoechst, blue). **D**. Quantification of (**C**) showing the percentage of SCs that had successfully associated and aligned to an axon. Each dot represents one independent experiment, data shown as mean ± SEM of 3 independent experiments. **E.** RT-qPCR analysis of RNA extracted from RafTR SCs treated with dbcAMP to induce differentiation, pre-treated with 0.5μM PD184352 (MEK-i) or solvent (DMSO), in the presence of absence of 10ng/ml TGF-β, for 30h. 4-hydroxytamoxifen (OH-Tmx) was used to activate the RafTR construct. **F.** RT-qPCR analysis of RNA extracted from SCs treated with dbcAMP to induce differentiation, transfected 48h prior to differentiation with 2nM Nf1-targeted (siNF1) or scrambled (scr) siRNA, in the presence of absence of TGF-β for 30h. Each dot represents one independent experiment. Data is normalised to B2m and shown as mean ± SEM of 3 independent experiments. For A and D-F, a two-way ANOVA was used. *p<0.05, **p<0.01, ***p<0.001, ****p<0.0001, ns=not significant. Not all statistical comparisons are shown.

**Supplementary Figure S8 – related to Figure 8.**
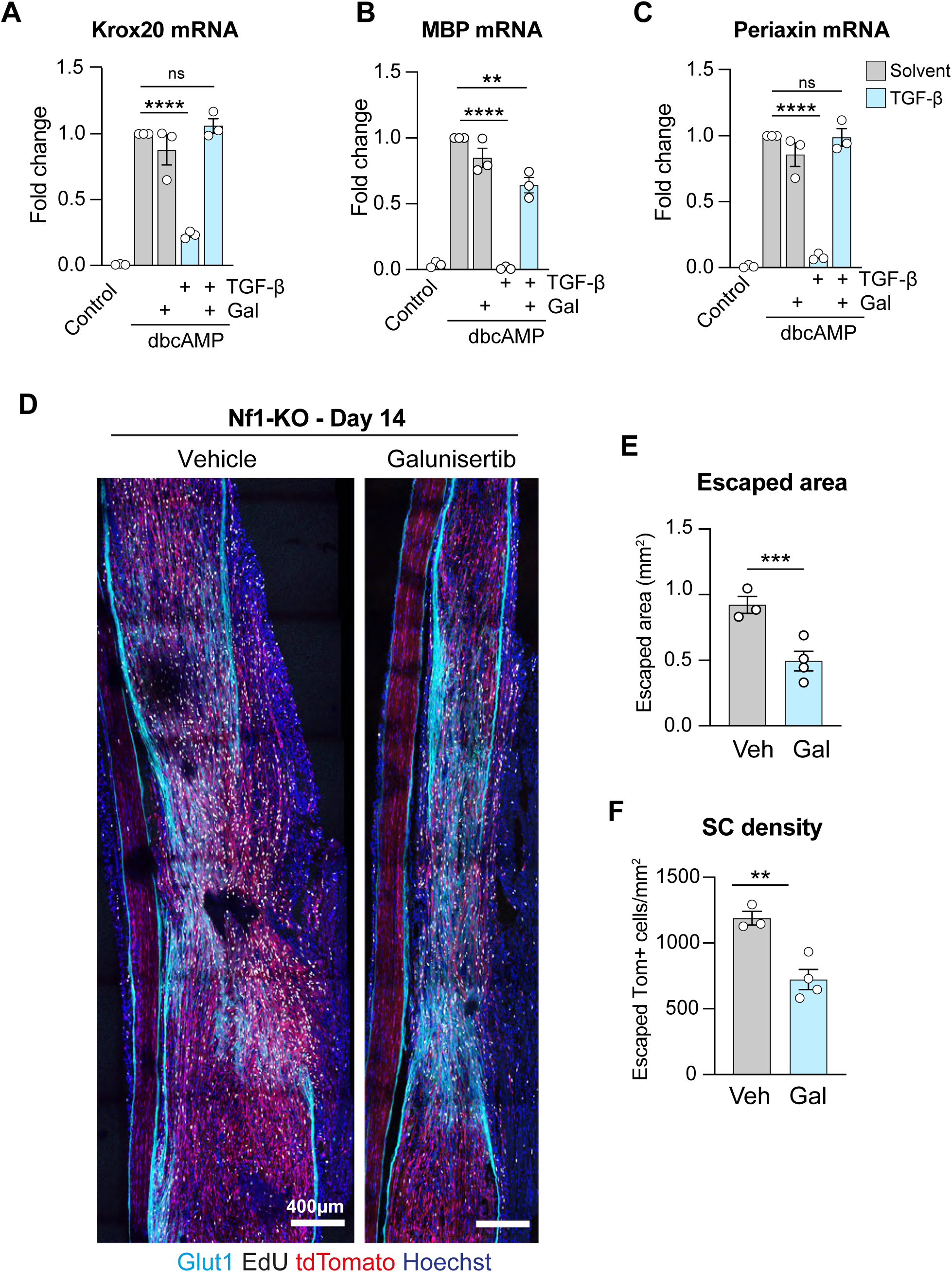
RT-qPCR analysis of RNA extracted from SCs treated with dbcAMP to induce differentiation, pre-treated with 10μM Galunisertib (Gal) or solvent (DMSO), in the presence of absence of 10ng/ml TGF-β, for 30h. (**A**) Krox20, (**B)** Myelin basic protein (MBP) and (**C**) Periaxin were used as indicators of differentiation. Each dot represents one independent experiment. Data is normalised to B2M levels and shown as mean ± SEM of 3 independent experiments. **D.** Representative tile scan confocal images of longitudinal sections of nerves from NF1-KO mice at Day 14 post-injury, treated with either vehicle or Galunisertib, twice a day at Days 7 – 9 after injury, showing recombined SCs (tdTomato, red), and labelled to detect EdU (white), perineurial cells (Glut1, cyan), and nuclei (Hoechst, blue). Quantification of (**D**) showing (**E**) the escaped area and (**F**) the escaped recombined SC (tdTomato+) density. Each dot represents one mouse (n=3-4). Data shown as mean ± SEM. For A-C, a two-way ANOVA was used; for E-F, an unpaired two-tailed t-test was used. **p<0.01, ***p<0.001, ****p<0.0001, ns=not significant. Not all statistical comparisons are shown.

## Notes

### Competing Interest Statement

The authors have declared no competing interest.

